# *Zfp423* regulates Sonic hedgehog signaling via primary cilium function

**DOI:** 10.1101/046961

**Authors:** Chen-Jei Hong, Bruce A. Hamilton

## Abstract

*Zfp423* encodes a 30-zinc finger transcription factor that intersects several canonical signaling pathways. *Zfp423* mutations result in ciliopathy-related phenotypes, including agenesis of the cerebellar vermis in mice and Joubert syndrome (JBTS19) and nephronophthisis (NPHP14) in humans. Unlike most ciliopathy genes, *Zfp423* encodes a nuclear protein and its developmental expression is complex, leading to alternative proposals for cellular mechanisms. Here we show that *Zfp423* is expressed by cerebellar granule cell precursors, that loss of *Zfp423* in these precursors leads to cell-intrinsic reduction in proliferation, loss of response to Shh, and primary cilia abnormalities that include diminished frequency of both Smoothened and IFT88 localization. Loss of *Zfp423* alters expression of several genes encoding key cilium components, including increased expression of *Tulp3*. *Tulp3* is a direct binding target of *Zfp423* and reducing the overexpression of *Tulp3* in *Zfp423*-deficient cells suppresses Smoothened translocation defects. These results define *Zfp423* deficiency as a bona fide ciliopathy, acting upstream of Shh signaling, and indicate a mechanism intrinsic to granule cell precursors for the resulting cerebellar hypoplasia.

**Author Summary:** Ciliopathies are a broad group of individually rare genetic disorders that share overlapping phenotypes and mutations in genes that make components of the primary cilium. Mutations in *ZNF423* are an exception. Patients and mouse models show characteristic hypoplasia of the cerebellar midline (Joubert syndrome), but the gene encodes a nuclear transcription factor. The mouse gene, *Zfp423*, is expressed in a dynamic developmental pattern, leaving the cellular mechanism for this brain malformation unresolved. One report suggested reduced Purkinje cell expression of Shh, a key mitogen for cerebellar granule cell precursors (GCPs) whose signal transduction occurs at the primary cilium, as the key event. We show that *Zfp423* mutants expressed normal Shh levels, but that *Zfp423*-depleted GCPs were unable to respond. Primary cilia on *Zfp423*-mutant GCPs in situ typically had a wider base and longer extension. ZNF423-depletion in a human cell culture model resulted in defective translocation of Smoothened, a key event in Shh signaling, and of the intraflagellar transport protein IFT88. RNA-Seq and RT-qPCR experiments identified known ciliopathy genes as potential conserved targets of *ZNF423* and *Zfp423*. One of these, *TULP3*, was both up-regulated in *ZNF423*/*Zfp423*-deficient cells and directly bound by Zfp423 in granule cell precursors. Reversing the overexpression of TULP3 in ZNF423-depleted human cell culture model reversed the defect in Smoothened translocation.

## Introduction

Cerebellar granule cell precursors (GCPs) are both an important model for neuronal development and a site of clinically important developmental abnormalities. GCP proliferation is highly responsive to Purkinje cell-derived sonic hedgehog (Shh) over a wide dynamic range [1–3]. Increasing or decreasing developmental Shh signaling can alter GCP dynamics sufficiently to reshape the cerebellum [4]. Correspondingly, ligand-independent signaling through the Shh pathway is a common feature in medulloblastoma, the most common pediatric brain tumor [5–7], and the Shh signaling pathway is a focal point of therapeutic development [8–10]. Shh signaling at primary cilia, where multiple components are trafficked to create a focused signaling module [11–14], is required for GCP expansion [15]. How Shh signaling is integrated with other signals that impact GCP proliferation, migration, and differentiation is not fully understood.

Through alternative interactions with multiple signaling and transcriptional pathways, Zfp423 (and its human ortholog, ZNF423) is well positioned to integrate extracellular signals into a coherent developmental response. Zfp423 was first described (as Roaz) through inhibitory interaction with EBF (Olf1) helix–loop–helix transcription factors [16]. Zfp423 is also a coactivator for BMP-activated SMADs [17, 18], retinoic acid receptors [19, 20], and Notch intracellular domain [21]. Intriguingly, Zfp423 activities on EBF and ligand-activated factors appear to be mutually inhibitory, further suggesting an integrative network function. *ZNF423* also appears to be a target of some cancers and low expression in neuroblastoma [20] or epigenetic silencing by Polycomb repressive complex 2 in glioma [22] is associated with poor prognosis.

Zfp423-deficient mice have a variety of developmental defects, including fully penetrant loss of cerebellar vermis [23–25] and variable loss of cerebellar hemispheres dependent on modifier genes and other factors [26]. Zfp423-deficient animals are also defective in forebrain development–including hypoplasia of the hippocampus and incomplete corpus callosum [23, 24], in olfactory neurogenesis [27], and in induction of adipose tissue [28, 29]. Mechanistically, literature to date has focused on physical interactions between Zfp423 and other transcription factors. Alternative levels of integration, such as alterations to cellular signaling centers, have not been well explored.

Hildebrandt and co-workers identified mutations in human *ZNF423* among patients with ciliopathy diagnoses [30]. Patients from all three *ZNF423* families reported had cerebellar vermis hypoplasia or Joubert Syndrome, while two also had nephronophthisis and other clinical features. Cellular assays with patient mutations showed effects on proliferation and DNA damage response, presenting a new pathogenic mechanism in ciliopathy disorders, but did not assess cilium structure or function. This raises the question of whether *ZNF423* and *Zfp423* mutations phenocopy ciliopathies by acting on downstream signaling events or represent bona fide ciliopathies by affecting cilium function upstream of signaling.

Results here provide new insights into Zfp423-dependent developmental mechanisms. Distinctly different models have been proposed for the cerebellar hypoplasia in *Zfp423* mice and, by extension, human patients. Based on *Zfp423* gene-trap expression in postnatal Purkinje cells, one group proposed a non-autonomous mechanism mediated by diminished Shh production [25], as seen in some Purkinje cell-selective mutations [31]. In situ hybridization, however, showed *Zfp423* expression in both the ventricular zone and the external germinal layer (EGL) in developing cerebellum, equally consistent with a GCP-intrinsic mechanism [23, 24]. We show that GCPs express Zfp423 protein in situ and in primary culture, that loss of Zfp423 blocked their ability to respond to Shh, altered cilium morphologies, decreased Smoothened translocation, and increased expression of several cilium-related genes, including *Tulp3*. Reversing the Tulp3 overexpression restored the frequency of Smoothened translocation. Our results demonstrate a GCP-intrinsic role for Zfp423 upstream of Shh signaling and suggest excess expression of target genes such as *Tulp3* as targets to improve function in ZNF423/Zfp423-deficient cells.

## Results

### *Zfp423* is expressed in granule cell precursors

In situ hybridization had previously shown *Zfp423* RNA expression in ventricular zone, external germinal layer and rhombic lip [23, 24], while lacZ reporter expression in a gene trap line suggested expression restricted to Purkinje cells [25], leading to different proposals for developmental defects in mutant embryos. To resolve which cells express Zfp423 in developing cerebellum, we examined Zfp423 protein expression and RNA in isolated cell populations (Fig 1). Zfp423 immunoreactivity showed strong, nuclear-limited signal in postmitotic Purkinje cells and most migrating GCPs in the rhombic lip and external germinal layer (EGL), with somewhat less intense staining in a fraction of ventricular zone cells (Fig 1A). Comparing sections from control (+/+) and *Zfp423* null mutant (*nur12*) congenic littermates that were processed in parallel on single slides demonstrated both specificity and sensitivity of the signal. Similar results have been obtained with two additional, independent antibodies developed against Zfp423 or its human ortholog (L. Flores-García and B.A.H.). As a further test of cell type distribution, we monitored by reverse transcription and quantitative polymerase chain reaction (RT-qPCR) the expression of *Zfp423* and several cell-type selective markers in cerebellar cells isolated by Percoll gradient centrifugation at E18.5 (Fig 1B,C). These data indicated that highly enriched GCP cultures retain *Zfp423* expression but not Purkinje cell selective markers such as *Rora* and *Shh*. Furthermore, RT-qPCR experiments showed similar RNA levels of *Shh*, the primary mitogen for GCPs, in *Zfp423*-mutant and control cerebellum among several independent samples (Fig 1D and S1_Table). These experiments demonstrate nuclear Zfp423 expression in GCPs in situ, with continued expression in primary culture, and support the potential for a cell-intrinsic role for Zfp423 in the granule cell deficits seen in *Zfp423*^−/−^ animals.

**Fig 1.**
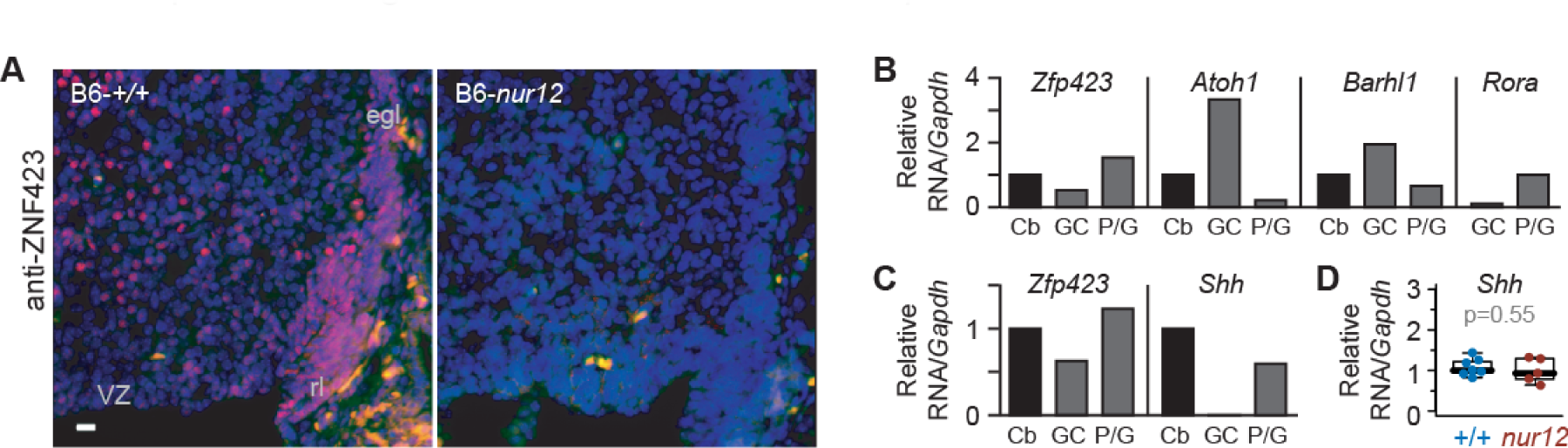
Zfp423 is expressed in cerebellar granule cell precursors. (***A***) Antibody staining at E16.5 shows nuclear expression of Zfp423 in cells of the rhombic lip (rl) and external germinal layer (egl), as well as ventricular zone (VZ) derivatives in control, but not in *Zfp423*^*nur12*^ mutant cerebellum. (***B***) E18.5 granule cell precursors purified by Percoll gradient centrifugation retain *Zfp423* expression. Quantitative RT-PCR shows co-expression of *Zfp423* with granule-specific *Atoh1* and *Barhl1* RNAs, with loss of Purkinje cell marker *Rora*. Histogram shows average expression ratio among technical replicates from a single purification, normalized to *Gapdh* and expressed relative to whole cerebellum. Cb, whole cerebellum; GC, granule cell/precursor fraction; P/G, Purkinje cell and Glial fraction. Rora level expressed relative to P/G fraction; Cb fraction not run. (***C***) Quantitative RT-PCR with hydrolysis probes (TaqMan) assays shows expression of *Zfp423* in the granule cell precursor fraction of a second independent purification. In contrast, no amplification from the granule cell preparation is detected for *Shh*, a GCP mitogen expressed by Purkinje cells. (***D***) RT-qPCR for *Shh* expression in whole cerebellum from control and *nur12* mutant animals dissected on P3. Mean values from technical replicates were normalized to *Gapdh* and expressed as a fraction of a control sample for 7 control and 5 mutant embryos. Similar results were obtained when normalized to *Ppia* cyclophilin. Group means were not significantly different (p=0.55, t-test).

### *Zfp423* is required for ex vivo proliferation and Shh responsiveness

To test whether GCP-intrinsic *Zfp423* expression is relevant to their proliferation phenotype, we transfected purified GCPs with shRNA vectors (Fig 2) that we previously validated for reducing Zfp423 levels [32]. Transfected GCPs were identified by fluorescence of an enhanced green fluorescent protein (EGFP) reporter in the vector. Cells entering S phase were marked by BrdU incorporation (A). The proportion of EGFP^+^ cells that were also BrdU^+^ was taken as a mitotic index specific for transfected cells. As predicted from *Zfp423* mutant animals [23], shRNA significantly reduced this index relative to the corresponding index of non-transfected (EGFP^−^) cells in the same cultures (p = 0.040-0.0014, paired *t* tests) and to approximately one-third the level of cells transfected by empty vector, irrelevant target (luciferase), or scrambled sequence shRNA controls (Fig 2B and S2_Table; p<10^−7^, one-factor ANOVA with post-hoc Tukey HSD test). Because the transfection efficiency in primary GCPs is modest (~15%), most transfected cells were surrounded by non-transfected cells. This minimizes the likelihood of any confounding effects secondary to disrupted cell interactions. These data show that *Zfp423* has a cell-intrinsic effect on GCP proliferation ex vivo and suggest that the effect might be cell autonomous within a population of GCPs, as proliferation phenotype does not appear to depend on the status of adjacent cells in the culture.

**Fig 2.**
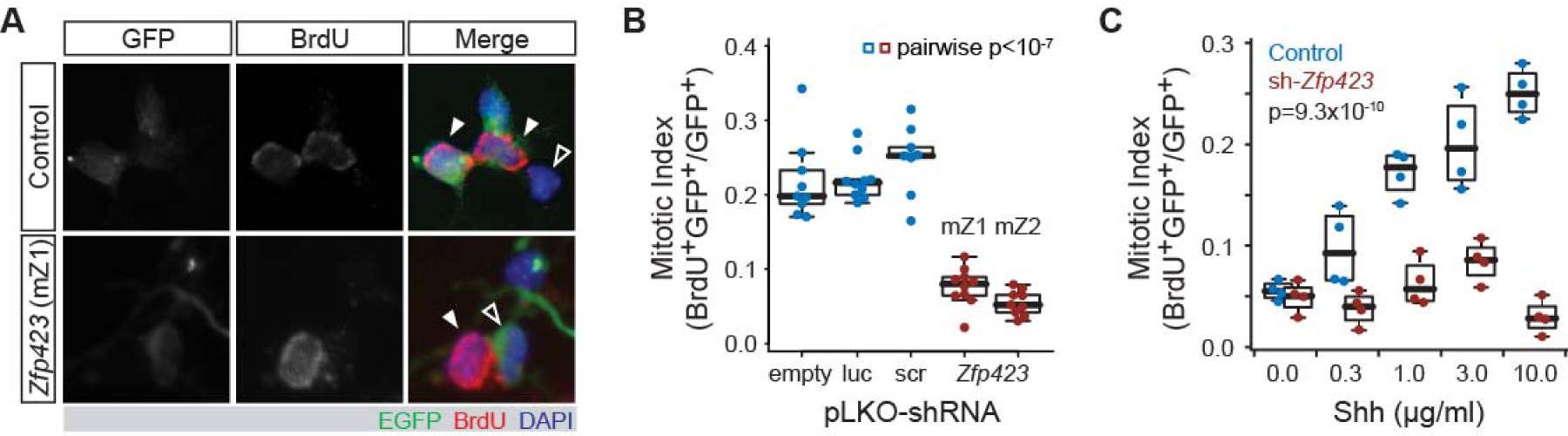
Zfp423 shRNA blocks proliferation and Shh response of purified granule cell precursors. (***A***) Examples of GCPs transfected with scrambled control or *Zfp423*-targeted shRNA constructs, in the presence of 3 µg/ml Shh. Transfected cells express EGFP from the shRNA vector (green). Proliferating cells are labeled by BrdU incorporation for 6 hr, at 48 hr after transfection (red). Control panels show dividing transfected cells (left-slanting closed arrowheads) and a non-dividing, non-transfected, cell (open arrowhead). *Zfp423* (shRNA mZ1) panels show a dividing non-transfected cell (right-slanting closed arrowhead) and a non-diving transfected cell (open arrowhead). At least two transfections for each of three independent DNA preparations were assayed for each construct. Channel levels were adjusted for merged images. (***B***) The proportion of EGFP^+^ cells labeled by BrdU is plotted for each replicate culture in the presence of 3 µg/ml Shh. shRNA sequence was a highly significant factor (p < 2.2 × 10^−16^, ANOVA), with each *Zfp423* shRNA construct different from every control shRNA (p < 10^−7^ for each pair-wise comparison, Tukey HSD after ANOVA). Controls were not significantly different from each other (p > 0.4), nor were the two *Zfp423* hairpin sequences (p > 0.4). Empty, pLKO vector with no hairpin sequence; luc, luciferase; scr, scrambled sequence control; mZ1 and mZ2 are non-overlapping shRNA sequences designed against *Zfp423* from the RNAi consortium collection. (***C***) GCP cultures were transfected with control (scr) or *Zfp423*-targeted (mZ1) shRNA constructs and treated with 0-10 µg/ml Shh. Transfected cells were recognized by EGFP fluorescence, replicating cells by BrdU incorporation. Plotted points indicate mitotic index of transfected cells for each of four replicate experiments at each of the indicated Shh concentrations. A minimum of three independent DNA preparations was used for transfection in each condition. The effect of *Zfp423*-targeting shRNA, Shh concentration, and the interaction between hairpin identity and Shh concentration were each strongly supported statistically (p= 9.3 × 10^−10^, 2.0 × 10^−4^, and 6.4 × 10^−6^ respectively, two-factor ANOVA).

Because previous reports showed that Shh is the principal mitogen for GCP proliferation in vivo and in culture, with a wide dynamic range, we next tested whether Zfp423-depletion in GCPs might alter the dose-response curve for exogenous Shh treatment (Fig 2C and S3_Table). We exposed Zfp423-knockdown and control GCPs to a range of 0-10 µg/ml recombinant Shh concentrations and measured the resulting mitotic index for replicate cultures at each concentration. Two-factor ANOVA on the full set of resulting cell count data showed significant effects of Shh dose (p = 2.0 × 10^−4^), Zfp423 vs. control shRNA treatment groups (p = 9.3 × 10^−10^), and the interaction between dose and shRNA (p = 6.4 × 10^−6^). Remarkably, Zfp423-shRNA cells were refractory to Shh levels more than an order of magnitude greater than that required to stimulate proliferation of control-shRNA cells. These results indicate a Zfp423-dependent step critical to Shh signal transduction in GCPs.

### Zfp423-deficient precursors have abnormal distribution of cilium morphology

Because Shh signaling is transduced through components localized to the primary cilium, we next asked whether Zfp423-deficient GCPs made cilia and basal bodies in normal frequency and quality (Fig 3). Cerebellum sections from E18.5 mice showed both structures intact among cells in the EGL and with comparable frequency per cell between *Zfp423*^*nur12*^-homozygous and littermate controls by double label immunofluorescence for acetylated α-tubulin (Ac-αTub) in the cilium and γ-tubulin (γTub) in the basal body (Fig 3A). Volume measurements extracted from optical sections, however, showed a significant increase in cilium volume in *Zfp423*^*nur12*^ EGL compared to littermate controls (Fig 3B and S4_Table). As replicate measurements did not follow a normal distribution within or between biological samples, we made several assessments using non-parametric tests. Among co-processed sections from eight littermate pairs, typical volumes were larger in the mutant than in the control animal for all eight comparisons (p = 0.0078, Wilcoxon signed rank test, two tailed). The distribution of volumes between genotypes for all 499 discrete measurements irrespective of pairing was also highly significant (p = 8.7 × 10^−6^, Kolmogorov-Smirnov test). Mutant cilia also had substantially higher variance than those in control littermates (0.89 vs. 0.33; p = 1.3 × 10^−4^, Ansari-Bradley test). Similar analysis of basal bodies (Fig 3C and S5_Table) did not support a difference between paired samples (p = 1), but did support a slightly lower distribution of basal body volumes over 662 individual measurements irrespective of pairs (p = 0.031) without significant difference in variances (p = 0.19). Mean voxel values for acetylated α-tubulin were 1.02 in control, 1.37 in mutant sections. Mean voxel values for γ-tubulin were 1.35 in control, 1.11 in mutant sections.

**Fig 3.**
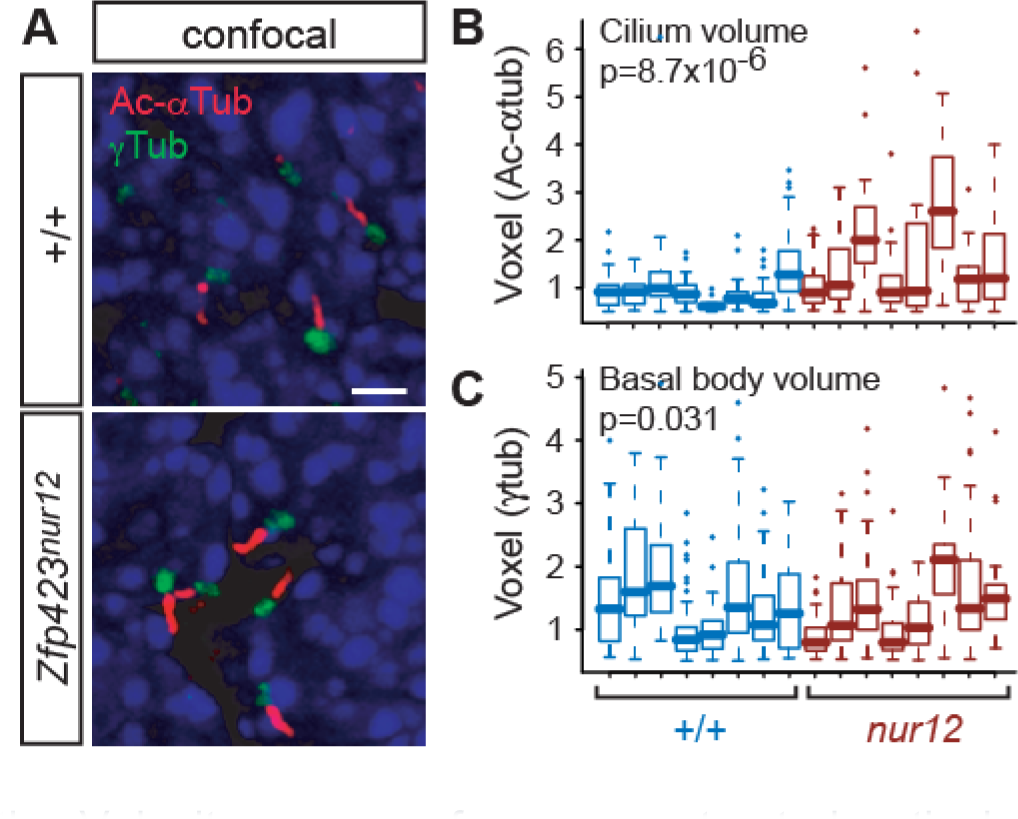
Confocal images show altered primary cilium volumes in Zfp423^nur12^ EGL. (***A***) Confocal micrographs show representative projection images showing cilia (anti-acetylated α-tubulin, Ac-αTub) and basal bodies (anti-γ-tubulin, γTub) in sections of E18.5 cerebellum from animals of the indicated *Zfp423* genotypes. All images are from sagittal sections, 0.7-0.9 mm from the midline. Scale bar, 2 µm. (***B,C***) Measured volumes for each of 499 axonemes (***B***) and 662 basal bodies (***C***) identified by the Volocity program from reconstructed optical sections. Measurements from paired mutant and control animals are color coded in the scatter plot. The differences in overall distributions between mutant and control are significant for both axonemes (p = 8.7 × 10^−6^) and basal bodies (p = 0.031, Kolmogorov-Smirnov test) in pooled data from eight pairs of animals.

To understand better the basis for these volume differences, we measured discrete parameters by structured illumination “super-resolution” microscopy of cilia from mutant and control littermate animals (Fig 4). We obtained similar results using either Ac-αTub (Fig 4A-C and S6_Table) or Arl13b (Fig 4D-F and S7_Table) as a marker protein. In Ac-αTub measurements, mutant cilia tended to be longer (Fig 4B), but this effect did not reach conventional statistical threshold (p = 0.079, Wilcoxon rank sum test, two tails), while mutant cilia had a significantly wider base compared with control (Fig 4C, p = 0.0091). Structured illumination measurements for Arl13b, associated with the ciliary membrane, showed significant increases in both length (Fig 4E, p=0.028) and base width (Fig 4F, p=0.013) of GCP cilia in an independent set of animals. Taken together, these observations indicate a cellular basis for ciliopathy-related phenotypes of *Zfp423*-deficient mice.

**Fig 4.**
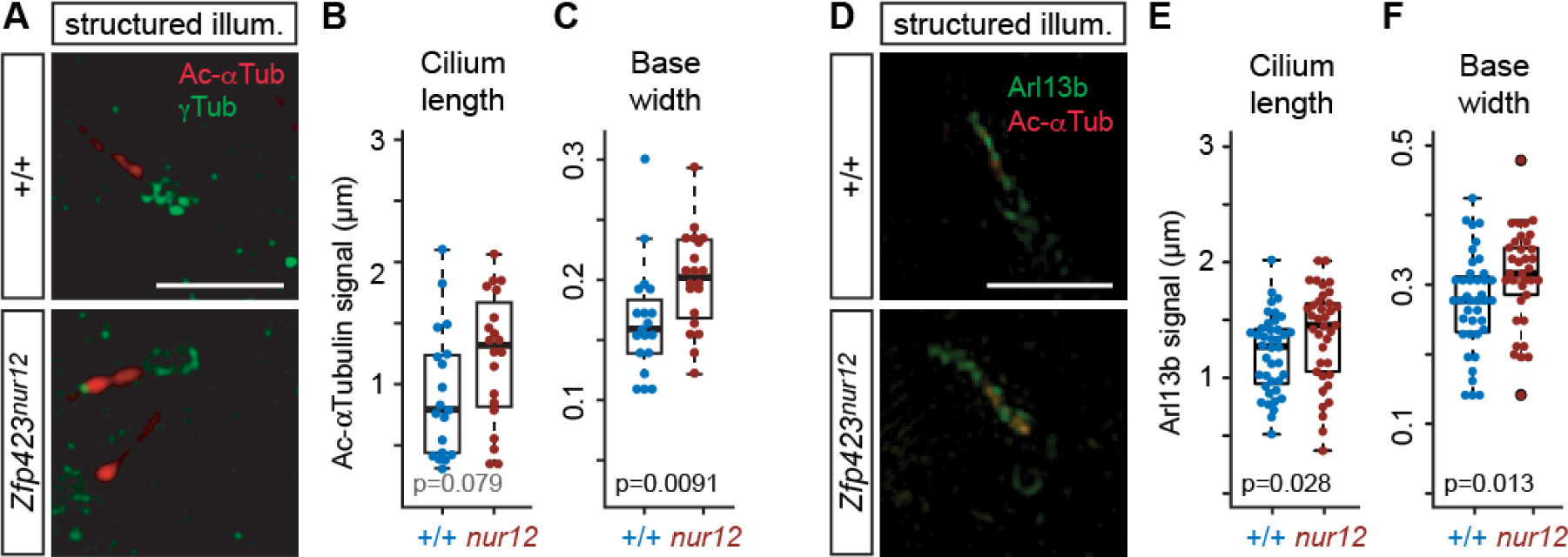
Structured illumination microscopy shows altered cilium dimensions in Zfp423^nur12^ EGL. (***A***) Super-resolution images show axoneme (Ac-αTub) and basal body (γTub). Scale bar, 2 µm. (***B,C***) Box and scatter plots show distributions of cilium length (***B***) and base width (***C***) measurements. Significance was assessed from paired samples. Length, p = 0.079, base width p = 0.0091, Wilcoxon rank sum test with two tails. (***D***) Structured illumination micrographs of cilia in the EGL. Arl13b and γ-tubulin in green, acetylated α-tubulin in red. (***E***) Length measurements from Arl13b signal in consecutively imaged cilia showed a shift toward greater length in *nur12* (p = 0.028, Wilcoxon rank sum test). (***F***) Width of the cilium base was also increased in *nur12* (p = 0.013, Wilcoxon rank sum test).

### *ZNF423* is required for quantitative Smoothened translocation during Shh signaling

Because Zfp423 is required for Shh response and affected the distribution of cilium morphologies, we asked whether it also affected trafficking of Smoothened, using a cell culture model (Fig 5). DAOY is a human medulloblastoma-derived cell line that expresses markers consistent with a GCP lineage [33, 34] and *ZNF423*, the human homologue of *Zfp423* (Fig 5A). DAOY cells were infected with a pseudotyped lentivirus expressing both a Smoothened-enhanced green fluorescent protein (EGFP) fusion protein and shRNA to either human *ZNF423* or a non-targeting control (Fig 5B). Cells were examined after serum starvation to produce a high percentage of cells with a definitive primary cilium. Two non-overlapping *ZNF423* shRNA were each effective in reducing its level of expression relative to multiple controls, a representative image is shown in Fig 5C. Quantification of fluorescence Western blots showed that, relative to control shRNAs, *ZNF423* knockdown cultures expressed ~30% ZNF423, ~110% acetylated α-tubulin, and ~95% IFT88 levels scaled to either GAPDH or β-actin as internal loading controls. We then tested the effect of ZNF423 knockdown on Smoothened localization to the cilium. Infected DAOY cells were treated with 0.1 µg/ml Shh as a low dose sufficient for signaling [1]. Cells were fixed 6 hr after stimulation and processed for immunofluorescence to visualize localization of the EGFP tag on Smoothened and acetylated α-tubulin as a marker of cilium location. Only EGFP^+^ cells were included to avoid measurements from uninfected cells. To compare localization of Smoothened-EGFP relative to acetylatedα-tubulin, immunofluorescence micrographs (Fig 5D) presented in randomized order were assessed categorically by five observers blinded to experimental conditions, for >800 cilia across five independent experiments. Summary results (Fig 5E and S8_Table) showed significantly less Smoothened-EGFP translocation in *ZNF423*-depleted cells (1.2×10^−13^, chi-square test). Results showed significant differences in each independent experiment (p = 0.0058 to 2.9×10^−5^).

**Fig 5.**
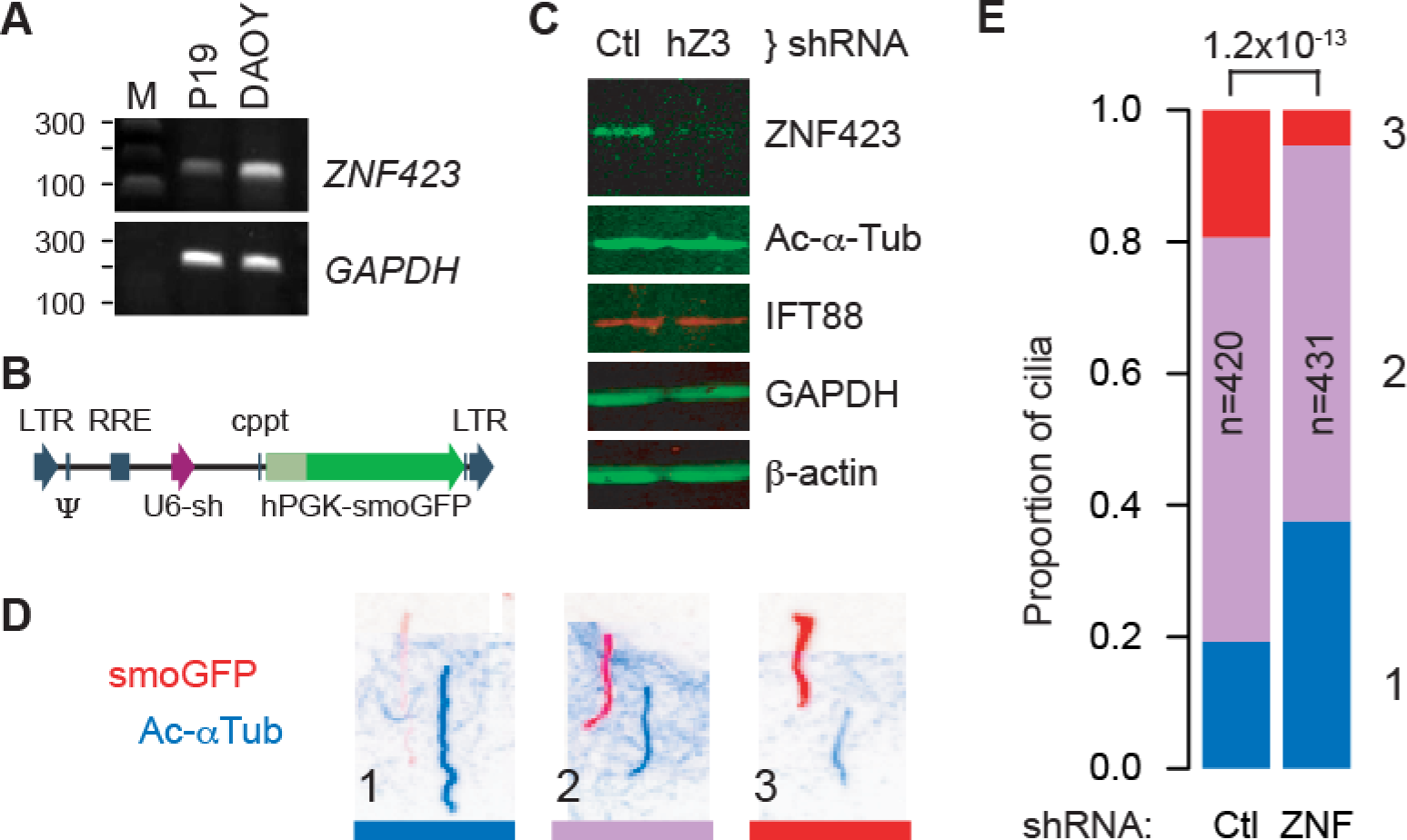
ZNF423 knockdown quantitatively alters Smoothened translocation. (***A***) RT-PCR shows *Zfp423* expression in mouse P19 cells [32] and *ZNF423* RNA expression in DAOY medulloblastoma cells. *GAPDH* was used as a positive control. M, size marker. (***B***) Structure of pLKO-derived lentivirus constructs, expressing a shRNA from a U6 promoter (purple arrow) and Smoothened-EGFP fusion (smoGFP, dark green) from a human *PGK* promoter (light green). Vector backbone elements are indicated in blue. (***C***) Western blots show reduced expression of ZNF423 protein in DAOY cells infected with lentivirus expressing a *ZNF423* shRNA (hZ3) relative to control (Ctl). (***D***) Color-inverted images of immunofluorescence for both GFP (red) and acetylated α-tubulin (blue) show Smoothened translocation phenotypes of cells infected with lentivirus expressing both smoGFP and an shRNA. Channel layers were shifted diagonally to allow independent view of each. Three categorical phenotypes were observed, (1) strong acetylated α-tubulin with little to no detected Smoothened, (2) intermediate levels, or (3) stronger Smoothened signal relative to acetylated α-tubulin. (***E***) Stacked histogram of cilium phenotypes summarizes five replicate experiments. Phenotypes are color-coded and numbered as in (D). Number of cilia scored in each group (n), *ZNF423*-directed (ZNF) or control (Ctl) shRNAs, and chi-square p-value for the difference between groups are indicated.

Mean fluorescence intensities along the length of each cilium were also calculated from the raw image files as a second analytical approach to the same experiments. Cilium annotations and measurements were performed blind to treatment group on images in randomized order. ZNF423-depleted cells showed ~30% reduction in both mean and median Smoothened-EGFP intensities in cilia, with a highly significant population shift between shRNA groups (p=7.0×10^−11^, Wilcoxon rank sum test; each of five replicates was independently significant, p<0.025). Ratios of Smoothened-EGFP to acetylated α-tubulin showed even stronger difference between groups (p<2.2×10^−16^), owing to an increase in acetylated α-tubulin signal in knockdown cells. These results strongly support a ZNF423-dependent step in Shh signaling upstream of Smoothened translocation in human DAOY cells.

### ZNF423 deficiency reduces IFT88 translocation in primary cilia

Because Smoothened is thought to use the intraflagellar transport (IFT) system for translocation into cilia [35, 36], we tested whether IFT88 localization was affected by loss of *ZNF423* in the DAOY culture model or *Zfp423* mutation in vivo (Fig 6). IFT88 is an essential component of the IFT-B complex required for quantitative hedgehog signaling in mice [11] and important for cerebellum development [37]. DAOY cells were infected with viruses described in Fig 5B and selected for cilium expression of the EGFP marker by direct fluorescence after fixation and staining. This necessarily restricted the analysis to the subpopulation of cells with strong Smoothened-EGFP translocation, as direct fluorescence is less sensitive than antibody staining and ZNF423-knockdown cells had a higher proportion of cilia near the detection limit. Among these cells, those transduced with *ZNF423*-directed shRNA showed both increased acetylated α-tubulin intensity relative to control (as predicted by Figures 4 and 5) and dramatically decreased ciliary IFT88 intensity (Fig 6A). The differences between *ZNF423* and control shRNAs reached very high statistical support for both mean intensities and ratios in cilia (Fig 6B-E and S9_Table, p<2.2x10^−16^, Kruskal-Wallis and Wilcoxon rank sum tests for full and pair-wise comparisons, respectively, in each measure), despite similar overall IFT88 protein levels in cellular extracts after *ZNF423* knockdown, as shown in Fig 5C. Strikingly, the IFT88 signal was also reduced relative to Smoothened-EGFP signal in *ZNF423*-knockdown cells compared to control (Fig 6E), suggesting that reduction in IFT88 staining was not a consequence of selection for the high Smoothened-EGFP population or comparison to acetylated α-tubulin levels.

**Fig 6.**
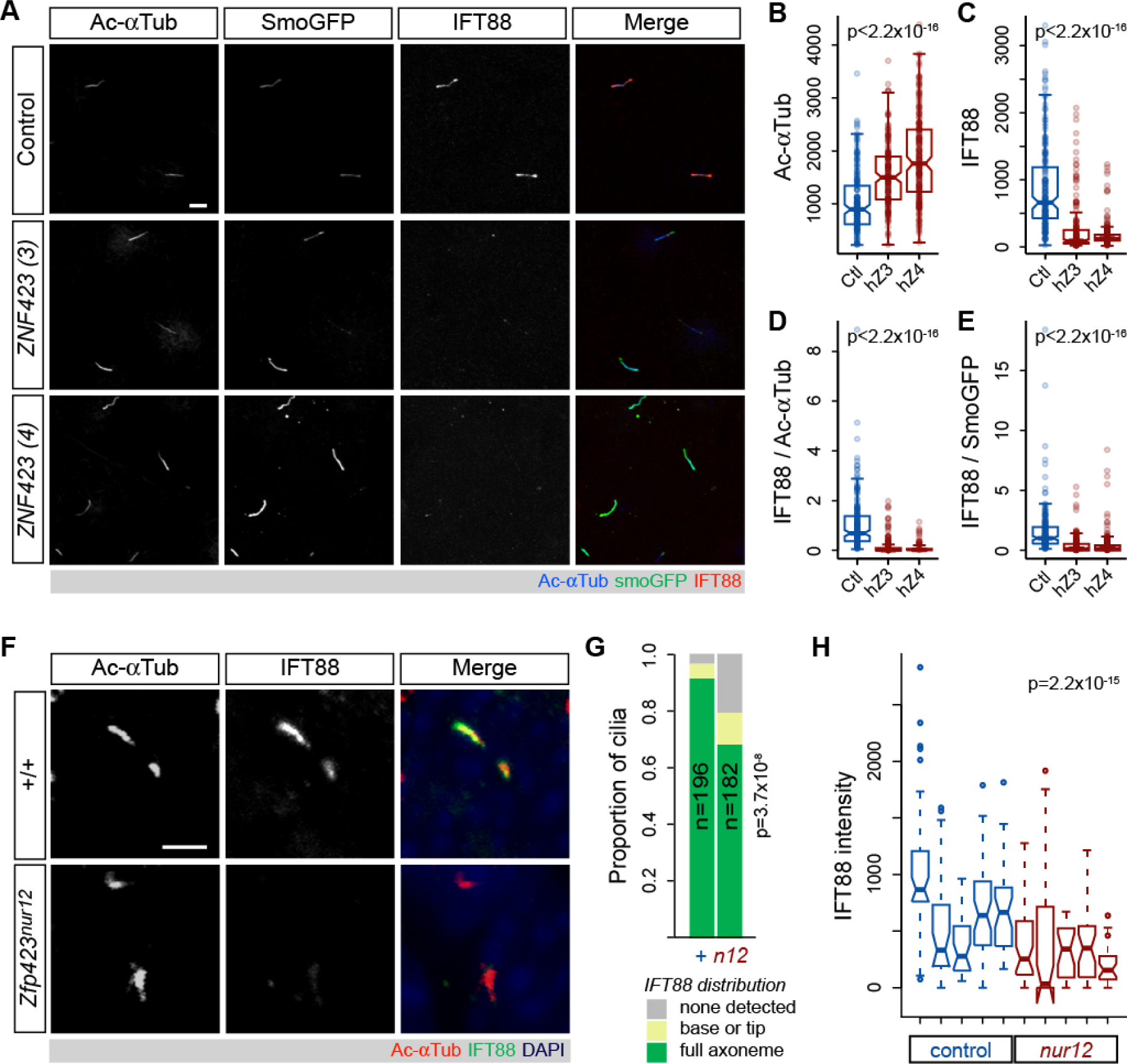
Reduced frequency and intensity of axonemal IFT88 in Zfp423-deificent cells. (***A***) Confocal micrographs of DAOY cells transduced with the indicated shRNA and Smoothened-EGFP. Immunofluorescence for acetylated α-tubulin and IFT88 and direct fluorescence of EGFP in representative images are shown. Scale bar, 5 µm. Distributions of mean axonemal measurements for (***B***) acetylated α-tubulin intensity, (***C***) IFT88 intensity, (***D***) ratio of IFT88 to acetylated α-tubulin and (***E***) ratio of IFT88 to Smoothened-EGFP fluorescence across three independent experiments with each shRNA in DAOY cells are plotted. For each plot, differences are highly significant among shRNA treatments (p<2.2×10^−16^, Kruskal-Wallis test) and between the 6 *ZNF423* and control populations (p<2.2×10^−16^, Wilcoxon rank sum test with continuity correction). (***F***) Confocal images of GCP cilia from EGL of control littermate and *Zfp423* null mutant (*nur12*) mice. Scale bar, 2 µm. (***G***) Localization of IFT88 relative to acetylated α-tubulin in cilia from five littermate pairs shows a lower proportion of mutant cilia with IFT88 throughout their length; p=3.7×10^−8^, chi-square test. (***H***) Mean axonemal intensities of IFT88 are significantly reduced compared to littermate controls. Differences between each littermate pair were independently significant. Difference between genotypes for summed data of 5 pairs is highly significant (p=2.2×10^−15^, Wilcoxon rank sum test).

To determine whether similar conditions affect granule precursor cells in situ, we examined the relative distributions of IFT88 and acetylated α-tubulin in the EGL of *Zfp423*^*nur12*^ and control littermate mice (Fig 6F-G). While most cilia from control littermates showed robust IFT88 staining in the cilium, many cilia in mutant EGL had little or no detectable IFT88 (Fig 6F). The proportion of cilia with little or no IFT88 staining was approximately three times greater among mutant animals than among their control littermates (Fig 6G and S10_Table). Similarly, the overall intensity of IFT88 within the cilium was significantly reduced in GCPs within the EGL of *nur12* animals compared with littermate controls (Fig 6H and S11_Table). Both measures showed highly significant *Zfp423*-dependence across five littermate pairs (p=3.7 × 10^−8^ for proportion, chi-square test; p=2.2 × 10^−15^ for signal intensities, Wilcoxon rank sum test with continuity correction). IFT88 translocation or retention thus appears impaired both in the human DAOY cell culture model and in mouse granule cell precursors in tissue sections, suggesting diminished opportunity to translocate Smoothened as a proximate cause for diminished Shh signaling both ex vivo and in situ.

### TULP3 overexpression is a functional target of *ZNF423* depletion

To link the functional requirement for human ZNF423 and mouse Zfp423 to potential target genes required for Shh response via cilia, we performed high-throughput cDNA sequencing (RNA-Seq) on the DAOY cell model, followed by confirmation of specific genes in freshly isolated mouse GCPs (Fig 7). RNA-Seq data were processed for normalized counts of annotated genes and then assessed for significant differences between *ZNF423* and control knockdown samples in three biological replicates, using tools in the HOMER software package [38, 39]. Gene set enrichment analysis [40, 41] did not identify any highly enriched pathways, but showed a strong relationship to genes up-regulated in DAOY cells after knockdown of *PCGF2*, a polycomb repressor complex protein [42]. Of >1000 genes that passed this initial threshold, 12 stood out for their known roles in primary cilia (Fig 7A). To determine whether these differences were conserved between the human medulloblastoma cell line model and mouse GCPs, we measured expression of mouse orthologs to five genes that had FDR < 0.01 and clean assays by RT-qPCR using freshly isolated GCPs from *Zfp423*^*nur12*^ and littermate controls (Fig 7B and S12_Table). *Tulp3*, *Nek9*, and *Arl4d* showed significantly elevated expression in mutant GCPs, as predicted by the DAOY cell knockdown model, while the other two were not significantly different between genotypes. In both DAOY and GCP cells, *TULP3*/*Tulp3* showed the most significant difference, being overexpressed >2-fold in each context.

**Fig 7.**
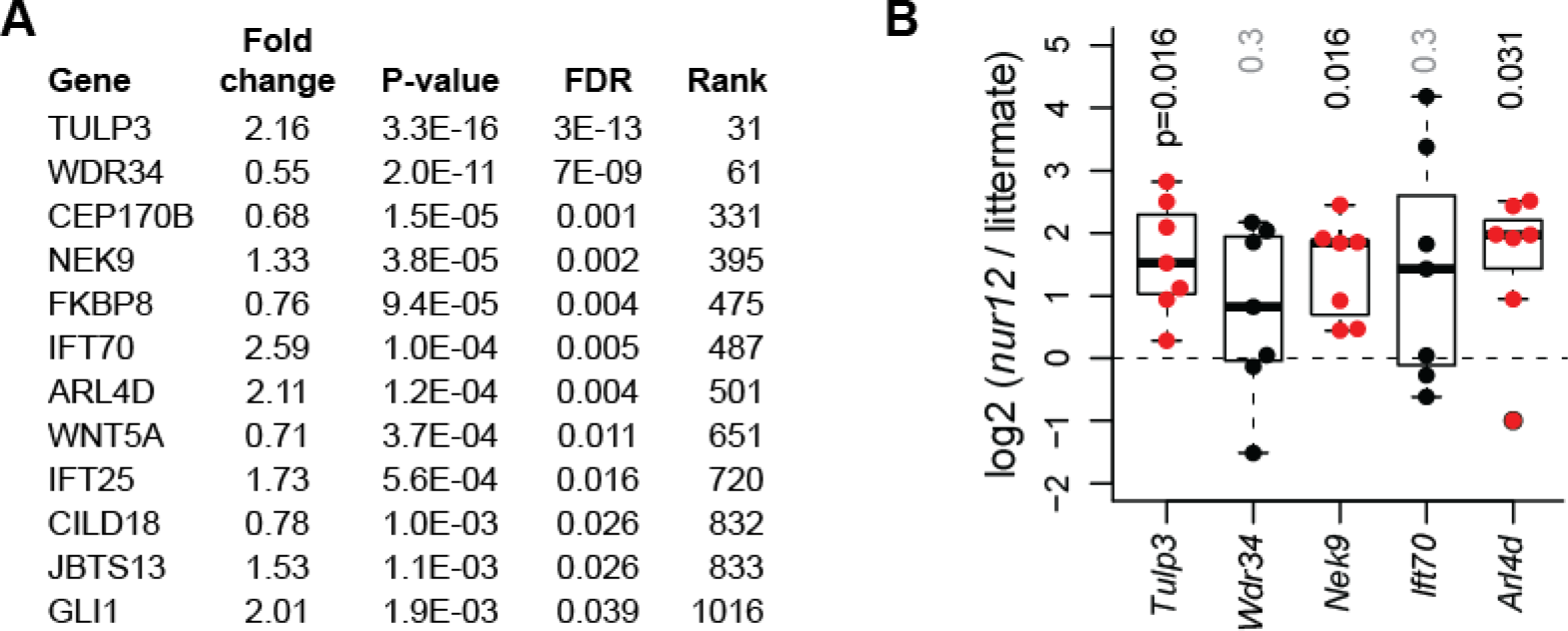
Tulp3 is an expression target of Zfp423 deficiency. (***A***) RNA-Seq data from knockdown and control DAOY cells identified several canonical cilium genes at high confidence. *TULP3* was among the largest magnitude changes and highest statistical confidence. (***B***) RT-qPCR from acutely isolated GCPs confirmed significantly increased expression of *Tulp3*, *Nek9*, and *Arl4d* in mutant relative to non-mutant controls among 7 littermate pairs (P0-P1). Nonparametric p-values from the Wilcoxon signed rank test are shown.

Chromatin immunoprecipitation (ChIP) experiments with independent antibodies demonstrated physical interaction of Zfp423 with the *Tulp3* gene in GCPs. Previous ChIP experiments quantified by high-throughput sequencing (ChIP-Seq) from a P19 cell culture model that expressed high levels of Zfp423, provided only modest read depth relative to cell number [32]. However, these data provided suggestive support for binding ~7 kb 5’ to *Arl4d* (Fig 8A) and for sites within and around *Tulp3* (Fig 8B). Because of the low yield in ChIP-Seq experiments and desire to test biological replicate samples, we quantified ChIP at these candidate sites relative to a previously described marker locus, 259C14S [43, 44], by ChIP-qPCR. Both a commercial antibody (goat) and a custom, affinity-purified antibody (rabbit) were previously characterized by Western blotting and immunofluorescence [32]. Both antibodies supported ChIP enrichment of a site in *Tulp3* intron 1 (Tulp3_3) that also had the strongest support in the previous ChIP-Seq data (Fig 8C and S13_Table). Although enrichment over background was modest with the commercial antibody, it was also uniquely significant at the Tulp3_3 site (p=0.037, Tukey Honest Significant Difference test after one-factor ANOVA). Enrichment with the custom antibody showed stronger support for the same site (p=0.012) and a non-significant trend at the Tulp3_2 site (p=0.28). Non-parametric analysis of pooled data from both antibodies supported the same result (p=0.0087, Wilcoxon rank sum test for Tulp3_3 vs. control site after Bonferroni correction, with no other pairwise combination p>0.2). Neither antibody showed enrichment for the other two candidate sites. Taken together, six independent ChIP experiments from primary GCPs support a direct and selective physical association of Zfp423 with sequences in *Tulp3* intron 1.

**Fig 8.**
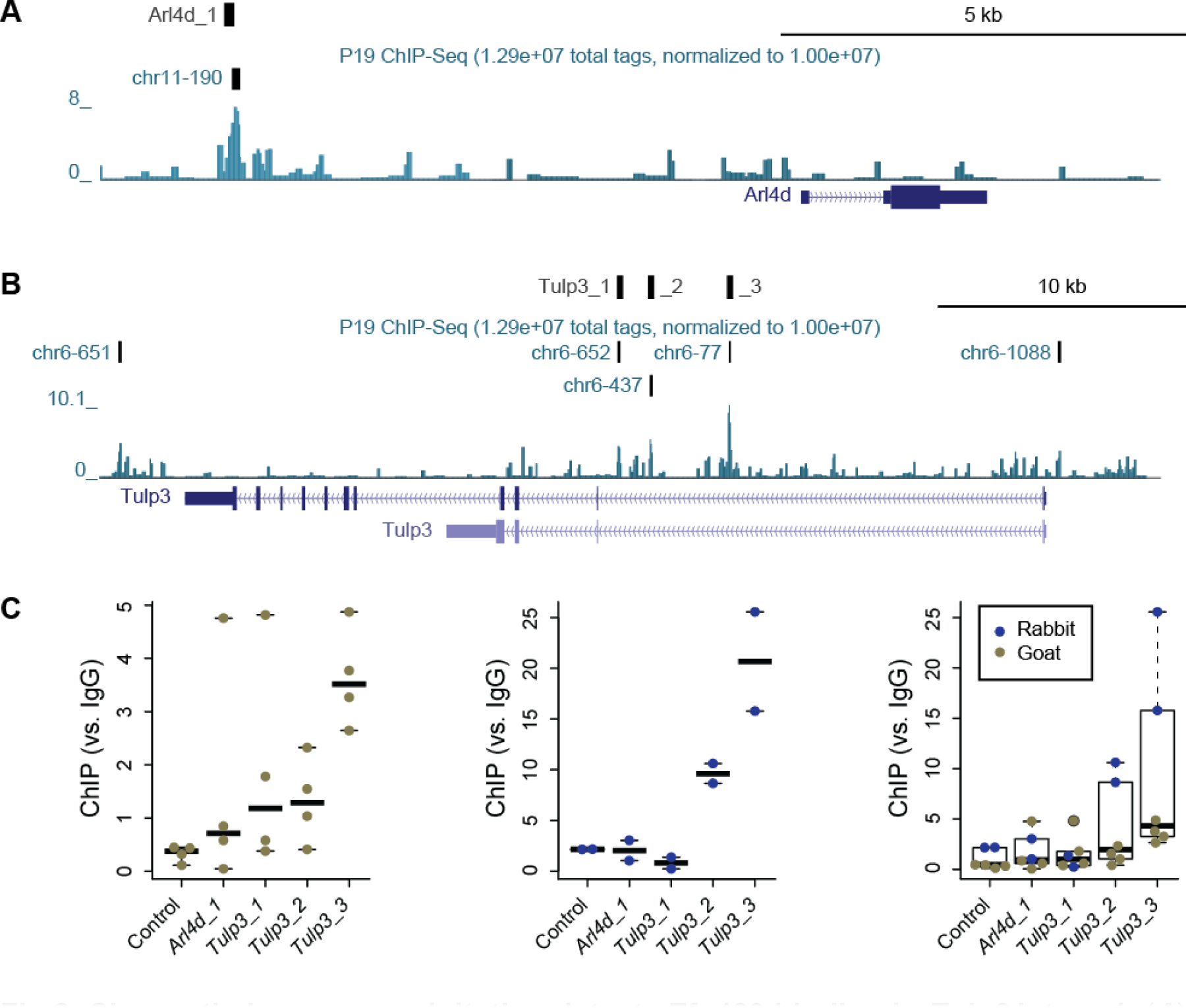
Chromatin immunoprecipitation detects Zfp423 binding in Tulp3 intron 1. (***A***) UCSC browser window on *Arl4d* locus shows location of the ChIP-PCR assay (Arl4d_1) and a ChIP-Seq peak call from P19 cells (chr11-190), 7 kb 5’ to the annotated transcript (Arl4d). The peak notation indicates the 190^th^ best-supported peak on chromosome 11 in that experiment [32]. (***B***) Similar window on *Tulp3* locus includes 5 potential peaks from P19 ChIP-Seq data, three of which supported reliable quantitative PCR assays (Tulp3_1, _2 and _3) within 150 bp of the peak. (***C***) Graphs show fold enrichment relative to an IgG control for a control locus and the four candidate binding sites. Colored dots indicate measured values from independent biological replicates, bars indicate median values for each locus. A commercial anti-Zfp423 antibody developed in goat (blue dots) showed modest but significant enrichment for Tulp3_3. A custom antibody developed in rabbit and affinity purified against the immunogen showed strong enrichment for Tulp3_3 and possible enrichment for Tulp3_2, 3 kb away. Non-parametric analysis of the combined data provides strong and conservative statistical support for binding at Tulp3_3 (p=0.0087, Wilcoxon rank sum test with Bonferroni correction).

To test the functional importance of TULP3 overexpression in effecting ZNF423 depletion phenotype, we compared knockdown of *ZNF423* to simultaneous knockdown of both *ZNF423* and *TULP3*, using the same approach and directly comparable to *ZNF423* knockdown in Fig 5. Combinations of targeting or control shRNAs were delivered to DAOY cells using lentiviral vectors. Knockdown of *ZNF423* with either of two non-overlapping shRNA depleted ZNF423 protein to ~50% of control levels (by Western blot) and produced ~2-fold increase in TULP3 protein in each culture (Fig 9A), even though only 60-75% cells appeared to be productively infected based on the fluorescent reporter protein. Of three shRNAs directed against *TULP3*, only T2 and T3 were effective in reducing measured TULP3 protein level (Fig 9B). T2 and T3 reduced TULP3 levels in *ZNF423*-knockdown cells to 70-90% of levels in control cells with full ZNF423 expression. We then compared Smoothened-EGFP translocation in doubly-infected cells, with ~100 cilia per group in each of two independent experiments (Fig 9C and S14_Table). Simultaneous knockdown of *TULP3* significantly reversed the reduction in Smoothened-EGFP translocation caused by knockdown *ZNF423* in each replicate experiment (p=0.0023 - 6.2×10-5, Pearson’s chi-squared test, 2 degrees of freedom for high vs. low TULP3). Each of the effective *TULP3* shRNAs showed similar result on Smoothened-EGFP distribution. These results indicate a functional role for TULP3 overexpression in mediating defective signaling in the primary cilium of ZNF423-deficient cells.

**Fig 9.**
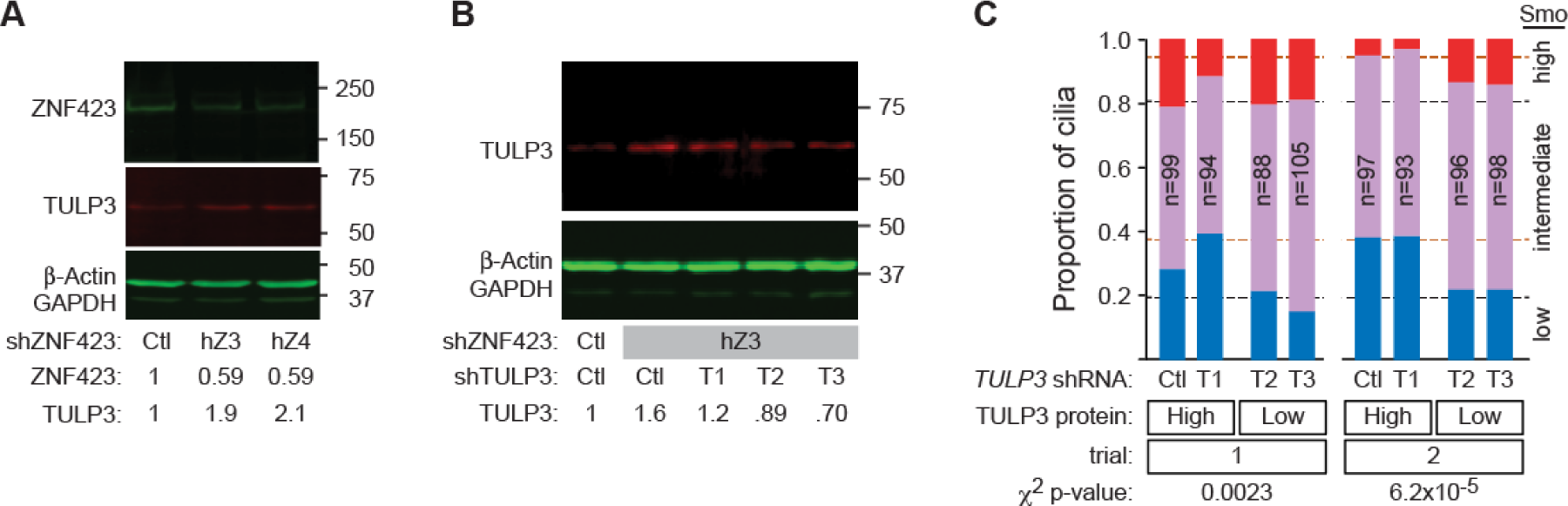
Tulp3 is a functional target of Zfp423. (***A***) Western blot shows loss of ZNF423 and increase of TULP3 levels in DAOY cells infected with lentiviral shRNA vectors targeting *ZNF423*. Relative amounts are compared to control shRNA infections after normalization to β-actin and GAPDH. (***B***) Western blot shows the effect of dual infection by *ZNF423* and three independent *TULP3* shRNA vectors on TULP3 protein. Only shRNAs T2 and T3 reduced TULP3 levels compared with control vector. (***C***) Co-infection with *ZNF423* and *TULP3* shRNA reverses the decrease in Shh-stimulated Smoothened GFP translocation into cilia seen with shRNA to *ZNF423* and either scrambled control or an ineffective *TULP3* shRNA. Graphs indicate high (red), medium (purple) and low (blue) translocation as in Fig 5D,E. Dashed lines indicate values from Fig 5E for *ZNF423* knockdown alone (orange dashes) and non-targeting controls (black dashes). Chi-square p-values are shown.

## Discussion

Our work provides, in both mouse brain and human cell culture models, the first mechanistic connection between ZNF423/Zfp423 ciliopathy-related phenotypes and function of the primary cilium. While phenotype severity and range of organ involvement vary, cerebellar vermis hypoplasia is a uniquely consistent feature among three reported *ZNF423* deficient patients [30] and five *Zfp423* mouse mutations on a variety of strain backgrounds [23–26]. Previous work in mice demonstrated loss of precursor proliferation [23], though the site of action has remained unresolved between competing proposals for cell intrinsic (potentially cell-autonomous) mechanisms in the granule cell lineage and non-autonomous mechanisms mediated by Shh expression in Purkinje cells. Analysis of cell culture models previously uncovered defective DNA damage response as a novel cellular mechanism in *ZNF423* patient-derived mutations [30], but did not assess effects on primary cilia and left open the question of whether ZNF423 might also act upstream of the cilium to regulate its function or downstream by mitigating its effects. Results presented here demonstrate a role for human and mouse homologs in cilium function and in intrinsic properties of granule cell precursors, acting – at least in part – through elevated expression of TULP3. In context, our work further unites cilium function and DNA damage response mechanisms by demonstrating that defects in both can occur through mutations for the same molecular component.

Four lines of evidence implicate *Zfp423* as a bona fide ciliopathy gene. First, both recessive loss of function and potentially dominant mutations produce phenotypes within the ciliopathy spectrum, including cerebellar vermis hypoplasia in both mice and humans, nephronophthisis in humans, and a variable degree of other consequences that may reflect allele strength or genetic, environmental, or stochastic modifiers [23, 24, 26, 30]. Second, our imaging analyses show that while *Zfp423*-null animals make primary cilia at a normal frequency, loss of Zfp423 results in an abnormal distribution of morphologies. As imaged with two marker proteins, cilia on mutant GCPs, as a group, appear wider near the base (Fig 3 and Fig 4), where several Joubert Syndrome and nephronophthisis proteins are localized [45–48], and tend to be longer. Structures at the base act to regulate traffic of signaling cargoes into and out of the cilium [49, 50]. This striking difference in cilium morphologies could be explained by an altered temporal profile of dynamic states, as the ranges of measurements largely overlap between genotypes despite having different distributions. Although it is possible that some portion of the shift in Ac-αTub-based measures could be due to changes in the local concentration of this modification in mice, knockdown of ZNF423 in the DAOY model does not dramatically change the overall level of Ac-αTub (Fig 5C). Additionally, any change in local Ac-αTub level would in itself indicate a functional change in cilium dynamics, as acetylation stabilizes the axonemal microtubules and de-acetylation is required for disassembly [51, 52]. Third, signaling through the primary cilium is functionally defective after loss of Zfp423 activity. *Zfp423*-depleted primary GCPs are refractory to exogenous Shh (Fig 2). *ZNF423*-depleted DAOY medulloblastoma cells are reproducibly deficient in translocation into the cilium of both Smoothened (Fig 5) and IFT88 (Fig 6), which are required for normal signal processing [12, 13, 53–55] and specifically for expansion of GCPs [15]. Diminished IFT88 translocation was also seen in mouse GCPs in situ (Fig 6), although the increased length distribution we observed is in contrast to the absence or severe shortening seen in Ift88 deficient cells [56, 57]. Fourth, we find expression level changes in key cilium components that are consistent between *ZNF423*-knockdown DAOY cells and *Zfp423*-mutant GCPs and indicative of disrupted signaling, at least one of which is functionally important and likely due to direct regulation. TULP3 was the best-supported target of Zfp423 in our analysis. TULP3 is an inhibitor of Shh signaling and loss of *Tulp3* in mice results in increased Shh signaling [58–60]. At a cellular level, TULP3 regulates ciliary trafficking at both retrograde transport and cilium entry through its interaction with IFT-A [61, 62]. ChIP experiments in primary GCPs with two different antibodies against Zfp423 both showed significant enrichment of a site in *Tulp3* intron 1 that was predicted from an earlier analysis in a P19 cell culture model [32]. Functionally, reversing the overexpression of TULP3 caused by ZNF423 depletion in the DAOY model significantly reversed the Smoothened translocation defects of *ZNF423* knockdown cells (Fig 9).

Although Zfp423 is required for mitogenic response to Shh ex vivo, residual proliferation both in knockdown cells and in the EGL of null animals away from the midline shows that Zfp423 is not strictly required for all mitogenic response by GCPs. Moreover, the DAOY model shows residual translocation of both IFT88 and Smoothened-EGFP in knockdown cells. One possibility might be that mitogenic response by GCPs is more sensitive to loss of *Zfp423* when cultured ex vivo than either GCPs in situ or the transformed DAOY model–and it is not unreasonable to expect that both environmental factors and intrinsic cell differences relevant to establishment of a cell line might enhance proliferative response. Alternatively (or additionally), it may be that IFT-independent diffusion of Smoothened into the cilium [50] provides a modest level of signaling that is saturated at lower levels of Shh than the translocation-dependent pathway, as might be suggested by the difference in sensitivity of Smoothened-EGFP and IFT88 localization to DAOY cell cilia (Fig 6E).

Tulp3 overexpression may be a surprising candidate for limiting Smoothened translocation. Although Tulp3 antagonizes Shh signaling, loss of *Tulp3* does not affect Smoothened trafficking [61, 63]. Rather, Tulp3 is thought to inhibit Shh signaling through localization of Gpr161, whose activation elevates cAMP level to promote cleavage of Gli proteins to their repressor forms [62]. However, this well-documented mechanism is based on Tulp3 loss-of-function may not be the only inhibitory effect when Tulp3 is expressed substantially above its normal level, as in ZNF423/Zfp423 deficient cells (Fig 7 and 9). Excess Tulp3 could in principle recruit additional inhibitory molecules to the cilium or titrate limiting components that promote Smoothened trafficking, such as Gprasp2 or Pifo [64, 65]. It is also possible that excess Tulp3 affects cell state in a way that indirectly affects IFT88 and Smoothened translocation to or concentration in the primary cilium. More work is needed to test such hypotheses, which are not mutually exclusive. In addition, since Zfp423 contributes to regulation of several ciliopathy-related genes, the full extent of molecular consequences is likely to be more complex and interplay among dysregulated components may be required to fully resolve molecular mechanisms behind the anatomical phenotypes seen in both patients and mouse models.

Our results have some similarities to those with ciliogenic transcription factors of the RFX family and Foxj1 [66, 67], but also important differences. RFX proteins are highly conserved regulators that activate core components for formation of motile cilia and loss of RFX activity results in loss or truncation of cilia [68–72]. *Foxj1* is a Shh target gene that also activates components to allow formation of motile cilia and is sufficient to drive their formation, with additional effects on primary cilia in some settings [73–76]. Restoring expression of target genes partially mitigates cell phenotypes [77]. By contrast, *ZNF423* patients and *Zfp423* mouse models present phenotypes more typical of defects in non-motile cilia, especially cerebellum vermis hypoplasia [23, 24, 30] and Zfp423 is not required for appearance of cilia. Zfp423 is required for efficient signaling through cilia in cerebellum and in the cell models we tested. Zfp423 appears to modulate components, often as a negative regulator, and we found improvement of one cellular phenotype by mitigating the overexpression of the best-supported repression target, *TULP3*, in a DAOY cell model.

The results here demonstrate a signaling mechanism, complementary to the DNA damage response (DDR) mechanism demonstrated by Chaki et al. [30], for neurodevelopmental phenotypes in *ZNF423*-deficient patients and orthologous *Zfp423* mutant mice. In addition to providing a structural and functional basis for diminished Shh signaling in granule cell precursors, our results point to loss of Zfp423 repressive functions on *Tulp3* and potentially other genes as a key component of the phenotype. Establishment of both DDR and signaling mechanisms provides a more comprehensive basis for understanding both the variety and variability of disease presentations in this ciliopathy, and potentially in others.

## Materials and Methods

*Mice.* The *nur12* mutation and BALB/c–*Zfp423*^*nur12*^ and C57BL/6–*Zfp423*^*nur12*^ congenic mice have been previously described [23, 26, 78]. For timed matings, midnight on the night of conception was considered to be E0.0.

### Antibodies

Anti-BrdU antibody (mouse monoclonal B44) was purchased from BD Biosciences. anti-GFP antibody (A11122) purchased from Molecular Probes. Antibodies against acetylated α-tubulin (mouse monoclonal 6-11B-1, T6793) and γ-tubulin (rabbit polyclonal, T5192) were purchased from Sigma-Aldrich. Rabbit anti-IFT88 (13967-1-AP), TULP3 (13637-1-AP), and Arl13b (17711-1-AP) were purchased from Proteintech. Mouse anti-GAPDH antibody (GT239) was purchased from GeneTex. Rabbit antiserum against Human ZNF423 (amino acid residues 247-407 relative to reference sequence NP_055884.2), was prepared and affinity-purified against the immunogen [32]. Commercial antibodies against ZNF423 (Santa Cruz Biotechnology sc-48785, aa 1-105; Sigma-Aldrich SAB2104426, aa 1234-1283) showed similar pattern (L. Flores-Garcia and B.A.H.). Western blots were developed with infrared-conjugated secondary antibodies (Rockland), detected on a Li-Cor Odyssey Imaging Station, and quantified in the ImageJ software package.

### Primary cell culture and mitotic index

Primary GCPs were isolated from E18.5 cerebella of BALB/c mice by Percoll gradient centrifugation essentially as described [79, 80], but omitting the adhesion step. Cells were transfected immediately after isolation and plated in the indicated concentration of Shh. A minimum of three independent DNA preparations was used for replication of measurements for each plasmid construct. 48 hr after transfection, cells were exposed to 4 µg/ml BrdU (Sigma-Aldrich) for 6 hr prior to fixation with 4% paraformaldehyde. Recombinant Shh was purchased from R&D Systems. TRC shRNAs 708 and 709 directed against mouse *Zfp423* were obtained from Sigma (Table 1).

**Table 1.**
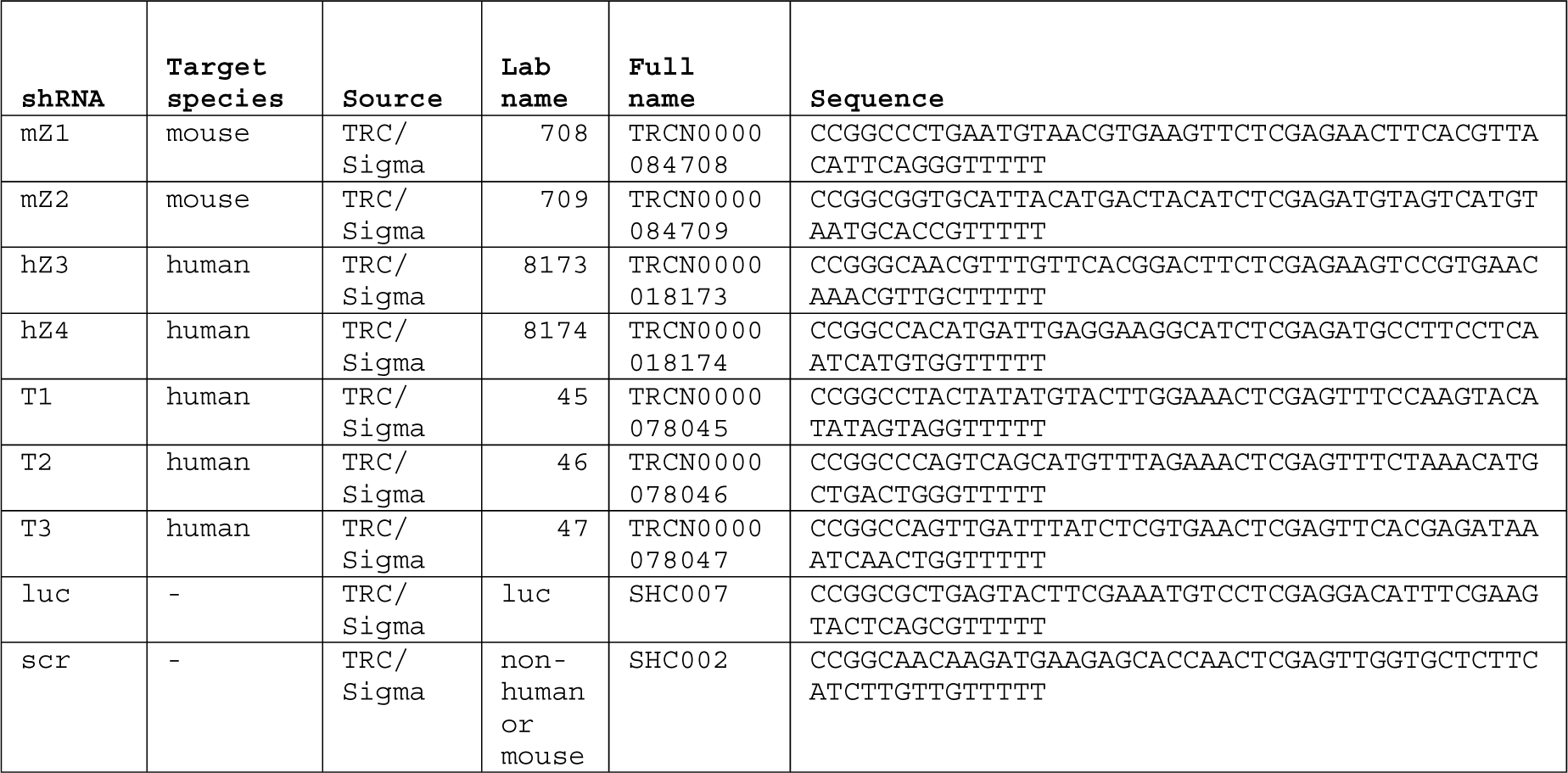
shRNA sequences.

### Gene expression assays

Quantitative RT-PCR was carried out on a Bio-Rad CFX-96 using SYBR Green fluorescence as described [43]. TaqMan hydrolysis probe assays (Life Technologies; *Shh* assay Mm00436528_m1, *Zfp423* assay Mm00473699_m1, *Gapdh* assay 4352339E) were performed on the same instrument, following manufacturer’s recommended conditions. Relative quantification used the ΔΔC_T_ method and comparisons across isolated cell fractions were expressed relative to the same ratio in whole cerebellum.

### Immunofluorescence and cilium measurements

For tissues, 10 µm sections were prepared from 4% paraformaldehyde fixed animals at age E18.5. Confocal (Olympus FV1000) and structured illumination “super resolution” microscopy (Applied Precision OMX) were performed in the UCSD School of Medicine Light Microscopy Facility. Volume measurements on the acquired images were calculated in Volocity software (PerkinElmer) from signal intensities across the full stack of images for each object, using the same intensity thresholds for co-processed mutant and control samples. For Smoothened translocation measurements, a Smoothened-EGFP fusion gene [81] was obtained from Addgene and cloned into pLKO.1 vector (Sigma), along with shRNA constructs targeting either *ZNF423*, scrambled control, or irrelevant sequence controls directed against reporter genes not used in this study. Human medulloblastoma-derived DAOY cells were obtained from Dr. Robert Wechsler-Reya. TRC shRNA directed against human *ZNF423* were obtained from Sigma (Table 1). Virus particles were packaged by co-transfection of 293FT cells with pCMVdR8.2 dvpr and pCMV-VSV-G (Addgene plasmid 8455 and 8454) as described [82]. Replicate cultures of DAOY cells were infected with each virus for 5 hr., cultured 7 days and serum starved for 24 hr prior to treatment with 0.1 µg/ml Shh for 24 hrs. Cells were fixed in 4% paraformaldehyde and imaged by confocal microscopy. Sixty-two to 100 consecutive images were collected for each sample and images from paired samples randomly ordered (knockdown vs. control) into a grid by the flip of a coin. Distribution of Smoothened-EGFP relative to acetylated α-tubulin was assessed as a nominal-scale variable by 5 different investigators blinded to genotype and experimental condition. Mean Intensity along the cilium was measured in each channel using ImageJ (National Institute of Health, USA).

### RNA-seq

ZNF423 knockdown and control DAOY cells were generated with a modified pLKO lentivirus that expressed shRNA to ZNF423 or a non-targeting control and co-expressed the fluorescent protein mCherry. Cells expressing high level of mCherry were enriched by flow cytometry and cultured in three replicate plates per virus. Bar-coded RNA libraries were prepared for sequencing using strand specific dUTP protocols [39] with minor modifications. Briefly, total RNA was harvested using Trizol reagent. Poly(A)^+^ RNA was enriched with Dynabeads mRNA Purification Kit (Life Technologies, Cat#61006). For each library, 500 ng poly(A)^+^ RNA was premixed with 0.2 μg of Oligo(dT) and random hexamer primer, heated to 70 °C for 10 min., and chilled on ice. Reverse transcription reactions were incubated at room temperature for 10 min., 42 °C 1h and 50 °C 20 min. First-strand cDNA was ethanol precipitated with 0.5 volumes 5 M ammonium acetate. After second-strand cDNA synthesis, sequencing libraries were constructed as described [38]. 250 to 500 bp size-selected, adaptor-ligated cDNA was incubated with 1 U USER (NEB) at 37 °C for 15 min. followed by 5 min. at 95 °C before PCR with Phusion High-Fidelity DNA polymerase. High-throughput sequencing was performed by UCSD IGM Genomics Center. Reads were aligned to the human hg19 transcriptome using STAR version 2.3.1f [83]. Data analysis was performed using HOMER [38] and the Bioconductor package edgeR [84, 85]. Gene set enrichment was performed in GSEA [40, 41]

### Chromatin Immunoprecipitation

ChIP was performed as described [32]. Briefly, 5-10 × 10^6^ purified primary GCPs were crosslinked in 1% formaldehyde, sonicated, and subjected to standard ChIP purification with the indicated antibodies at ~2 µg. Quantitative PCR from ChIP samples used 0.5% input fraction and ChIP with pre-immune IgG for relative quantification among samples.

### Statistical tests

Standard statistical procedures were performed in the R base environment (R 2.8.1 32-bit (5301) or R 3.0.3 64-bit (6660)). Normality (Shapiro-Wilk test) and equal variance (F test) of each data set was assessed prior to selecting parametric (ANOVA, *t*) or non-parametric (Kruskal-Wallis, Wilcoxon) tests. Tukey box-and-whisker plots indicate median (heavy line), quartile (box), and 1.5 times interquartile range (whiskers). Notches where shown indicate ±1.58 times the interquartile range/square root of *n*.

### Ethics statement

Mice were euthanized by CO2 inhalation or by perfusion or organ removal under deep anesthesia with tribromoethanol (avertin). All vertebrate animal procedures were approved by the University of California San Diego Institutional Animal Care and Use Committee (IACUC). The University of California San Diego is AAALAC accredited, AAALAC institutional number 000503.

## Accession Numbers

RNA-seq data have been submitted and can be accessed by the Gene Expression Omnibus (GEO) accession number GSE59598.

## Acknowledgements

The authors gratefully acknowledge Robert Wechsler-Reya for DAOY cells, Young-Wook Cho for affinity-purified ZNF423 antibody, Wendy Alcaraz, Dorothy Concepcion, Lisbeth Flores-Garcia, Kevin Ross, Isabella Sanchez, Weica Xie, and Susan Zhang for manual scoring of de-identified images, Jennifer Santini of the UCSD Neuroscience Microscopy Shared Facility for advice and assistance with instrumentation, Sven Heinz and Christopher K. Glass for advice on RNA-Seq library preparation and HOMER software, the UCSD Institute for Genomic Medicine Genomics Core for high-throughput sequencing and Lawrence S. B. Goldstein and Xiang-Dong Fu for comments on draft manuscripts.

## Supporting Information

S1-S14 Tables provide machine-readable data for the graphical figure panels with which they are referenced in the text.

**S1 Table. RT-qPCR data for Fig 1D.**

RT-qPCR measures and technical replicate standard deviations for *Shh* RNA relative to either *Gapdh* or *Ppia* internal control standards are shown for each sample.

**S2 Table. Mitotic index data for Fig 2B.**

For each shRNA, three independent DNA preparations were used in multiple transfections as replicate samples. Numbers of EGFP and BrdU double-positive, EGFP positive BrdU negative, EGFP negative BrdU positive, and double negative cells scored in each transfection is given.

**S3 Table. Mitotic index data for Fig 2C.**

For each indicated Shh (µg/ml), shRNA, DNA preparation, and transfection experiment, EGFP positive (transfected) cells were counted as BrdU positive (Brdu) or BrdU unlabeled (Un).

**S4 Table. Acetylated α-tubulin volume measures for Fig 3B.**

Columns indicate the littermate pair, animal, *nur12* (n) or wild-type (w) genotype and cilium voxel measurement from acetylated α-tubulin immunofluorescence.

**S5 Table. Basal body γ-tubulin volume measures for Fig 3C.**

Columns indicate the littermate pair, animal, *nur12* (n) or wild-type (w) genotype and basal body voxel measurement from γ-tubulin immunofluorescence.

**S6 Table. Structured illumination measures of cilium acetylated α-tubulin for Fig 4B-C.**

Genotype of *nur12* (n12) or +/+ littermate (wt), measured cilium length and base width are tabulated as graphed in Fig 4B and C.

**S7 Table. Structured illumination measures of cilium Arl13b for Fig 4E-F.**

Genotype of *nur12* (nur12) or +/+ littermate (control), measured cilium length and base width are tabulated as graphed in Fig 4E and F.

**S8 Table. Smoothened-EGFP translocation scores Fig 5E.**

For five independent trials the index image, shRNA to ZNF423 (n) or control (wt), and the median of categorical scores (Fig 5D) given by independent reviewers are tabulated.

**S9 Table. Immunofluorescence measures for Fig 6B-E.**

For three independent trials, the shRNA and its target (and an overall classifier for target, shRNA and trial number) are given with image values for IFT88, Smoothened-EGFP, and acetylated α-tubulin for each cilium.

**S10 Table. Ift88 distribution data for Fig 6G.**

For paired littermates, the pair number, *Zfp423* genotype (wt or *nur12* homozygous) and frequency of Ift88 staining at tips only, throughout the full cilium, or not detected (nd) and total number of cilia imaged for each animal are given.

**S11 Table. Ift88 intensity data for Fig 6H.**

Littermate pair, *Zfp423* genotype, and Ift88 intensity measures are shown.

**S12 Table. Paired RT-qPCR measures for Fig 7B.**

For paired littermate tissue samples, normalized RT-qPCR values are given for the *Zfp423*^*nur12*^ mutant and its control littermate for each of the five genes tested.

**S13 Table. ChIP-qPCR data for Fig 8C.**

For each of five indicated PCR assays, the antibody used, experimental trial, Cq and ΔCq values, and measured level normalized to input and IgG control are provided.

**S14 Table. Smoothened-EGFP translocation scores for Fig 9C.**

For doubly transfected cells, each shRNA virus used, the categorical translocation score assigned by five independent raters along with the median and mean scores are given.

## References

1. Wechsler-Reya RJ, Scott MP. Control of neuronal precursor proliferation in the cerebellum by Sonic Hedgehog. Neuron. 1999;22(1):103–14. PubMed PMID: 10027293.

2. Wallace VA. Purkinje-cell-derived Sonic hedgehog regulates granule neuron precursor cell proliferation in the developing mouse cerebellum. Curr Biol. 1999;9(8):445–8. PubMed PMID: 10226030.

3. Dahmane N, Ruiz-i-Altaba A. Sonic hedgehog regulates the growth and patterning of the cerebellum. Development. 1999;126(14):3089–100. PubMed PMID: 10375501.

4. Corrales JD, Blaess S, Mahoney EM, Joyner AL. The level of sonic hedgehog signaling regulates the complexity of cerebellar foliation. Development. 2006;133(9):1811–21. Epub 2006/03/31. doi: 10.1242/dev.02351. PubMed PMID: 16571625.

5. Marino S. Medulloblastoma: developmental mechanisms out of control. Trends Mol Med. 2005;11(1):17–22. Epub 2005/01/15. doi: 10.1016/j.molmed.2004.11.008. PubMed PMID: 15649818.

6. Roussel MF, Hatten ME. Cerebellum development and medulloblastoma. Curr Top Dev Biol. 2011;94:235–82. Epub 2011/02/08. doi: 10.1016/B978-0-12-380916-2.00008-5. PubMed PMID: 21295689.

7. Wechsler-Reya R, Scott MP. The developmental biology of brain tumors. Annu Rev Neurosci. 2001;24:385–428. PubMed PMID: 11283316.

8. Stanton BZ, Peng LF. Small-molecule modulators of the Sonic Hedgehog signaling pathway. Mol Biosyst. 2010;6(1):44–54. Epub 2009/12/22. doi: 10.1039/b910196a. PubMed PMID: 20024066.

9. Hyman JM, Firestone AJ, Heine VM, Zhao Y, Ocasio CA, Han K, et al. Small-molecule inhibitors reveal multiple strategies for Hedgehog pathway blockade. Proc Natl Acad Sci U S A. 2009;106(33):14132–7. Epub 2009/08/12. doi: 10.1073/pnas.0907134106. PubMed PMID: 19666565; PubMed Central PMCID: PMC2721821.

10. Romer J, Curran T. Targeting medulloblastoma: small-molecule inhibitors of the Sonic Hedgehog pathway as potential cancer therapeutics. Cancer Res. 2005;65(12):4975–8. Epub 2005/06/17. doi: 10.1158/0008-5472.CAN-05-0481. PubMed PMID: 15958535.

11. Huangfu D, Liu A, Rakeman AS, Murcia NS, Niswander L, Anderson KV. Hedgehog signalling in the mouse requires intraflagellar transport proteins. Nature. 2003;426(6962):83–7. Epub 2003/11/07. doi: 10.1038/nature02061. PubMed PMID: 14603322.

12. Corbit KC, Aanstad P, Singla V, Norman AR, Stainier DY, Reiter JF. Vertebrate Smoothened functions at the primary cilium. Nature. 2005;437(7061):1018–21. Epub 2005/09/02. doi: 10.1038/nature04117. PubMed PMID: 16136078.

13. Haycraft CJ, Banizs B, Aydin-Son Y, Zhang Q, Michaud EJ, Yoder BK. Gli2 and Gli3 localize to cilia and require the intraflagellar transport protein polaris for processing and function. PLoS Genet. 2005;1(4):e53. Epub 2005/10/29. doi: 10.1371/journal.pgen.0010053. PubMed PMID: 16254602; PubMed Central PMCID: PMC1270009.

14. Huangfu D, Anderson KV. Cilia and Hedgehog responsiveness in the mouse. Proc Natl Acad Sci U S A. 2005;102(32):11325–30. Epub 2005/08/03. doi: 10.1073/pnas.0505328102. PubMed PMID: 16061793; PubMed Central PMCID: PMC1183606.

15. Spassky N, Han YG, Aguilar A, Strehl L, Besse L, Laclef C, et al. Primary cilia are required for cerebellar development and Shh-dependent expansion of progenitor pool. Dev Biol. 2008;317(1):246–59. Epub 2008/03/21. doi: 10.1016/j.ydbio.2008.02.026. PubMed PMID: 18353302.

16. Tsai RY, Reed RR. Cloning and functional characterization of Roaz, a zinc finger protein that interacts with O/E-1 to regulate gene expression: implications for olfactory neuronal development. J Neurosci. 1997;17(11):4159–69. PubMed PMID: 9151733.

17. Hata A, Seoane J, Lagna G, Montalvo E, Hemmati-Brivanlou A, Massague J. OAZ uses distinct DNA- and protein-binding zinc fingers in separate BMP-Smad and Olf signaling pathways. Cell. 2000;100(2):229–40. PubMed PMID: 10660046.

18. Ku MC, Stewart S, Hata A. Poly(ADP-ribose) polymerase 1 interacts with OAZ and regulates BMP-target genes. Biochem Biophys Res Commun. 2003;311(3):702–7. PubMed PMID: 14623329.

19. Holzel M, Huang S, Koster J, Ora I, Lakeman A, Caron H, et al. NF1 is a tumor suppressor in neuroblastoma that determines retinoic acid response and disease outcome. Cell. 2010;142(2):218–29. Epub 2010/07/27. doi: 10.1016/j.cell.2010.06.004. PubMed PMID: 20655465; PubMed Central PMCID: PMC2913027.

20. Huang S, Laoukili J, Epping MT, Koster J, Holzel M, Westerman BA, et al. ZNF423 is critically required for retinoic acid-induced differentiation and is a marker of neuroblastoma outcome. Cancer Cell. 2009;15(4):328–40. PubMed PMID: 19345331.

21. Masserdotti G, Badaloni A, Green YS, Croci L, Barili V, Bergamini G, et al. ZFP423 coordinates Notch and bone morphogenetic protein signaling, selectively up-regulating Hes5 gene expression. J Biol Chem. 2010;285(40):30814–24. PubMed PMID: 20547764.

22. Signaroldi E, Laise P, Cristofanon S, Brancaccio A, Reisoli E, Atashpaz S, et al. Polycomb dysregulation in gliomagenesis targets a Zfp423-dependent differentiation network. Nat Commun. 2016;7:10753. Epub 2016/03/01. doi: 10.1038/ncomms10753. PubMed PMID: 26923714; PubMed Central PMCID: PMC4773478.

23. Alcaraz WA, Gold DA, Raponi E, Gent PM, Concepcion D, Hamilton BA. Zfp423 controls proliferation and differentiation of neural precursors in cerebellar vermis formation. Proc Natl Acad Sci U S A. 2006;103(51):19424–9. PubMed PMID: 17151198.

24. Cheng LE, Zhang J, Reed RR. The transcription factor Zfp423/OAZ is required for cerebellar development and CNS midline patterning. Dev Biol. 2007;307(1):43–52. PubMed PMID: 17524391.

25. Warming S, Rachel RA, Jenkins NA, Copeland NG. Zfp423 is required for normal cerebellar development. Mol Cell Biol. 2006;26(18):6913–22. PubMed PMID: 16943432.

26. Alcaraz WA, Chen E, Valdes P, Kim E, Lo YH, Vo J, et al. Modifier genes and non-genetic factors reshape anatomical deficits in Zfp423-deficient mice. Hum Mol Genet. 2011;20(19):3822–30. Epub 2011/07/07. doi: 10.1093/hmg/ddr300. PubMed PMID: 21729880; PubMed Central PMCID: PMC3168291.

27. Cheng LE, Reed RR. Zfp423/OAZ participates in a developmental switch during olfactory neurogenesis. Neuron. 2007;54(4):547–57. PubMed PMID: 17521568.

28. Gupta RK, Arany Z, Seale P, Mepani RJ, Ye L, Conroe HM, et al. Transcriptional control of preadipocyte determination by Zfp423. Nature. 2010;464(7288):619–23. PubMed PMID: 20200519.

29. Zhang LJ, Guerrero-Juarez CF, Hata T, Bapat SP, Ramos R, Plikus MV, et al. Innate immunity. Dermal adipocytes protect against invasive Staphylococcus aureus skin infection. Science. 2015;347(6217):67–71. Epub 2015/01/03. doi: 10.1126/science.1260972. PubMed PMID: 25554785; PubMed Central PMCID: PMC4318537.

30. Chaki M, Airik R, Ghosh AK, Giles RH, Chen R, Slaats GG, et al. Exome Capture Reveals ZNF423 and CEP164 Mutations, Linking Renal Ciliopathies to DNA Damage Response Signaling. Cell. 2012;150(3):533–48. Epub 2012/08/07. doi: 10.1016/j.cell.2012.06.028. PubMed PMID: 22863007.

31. Gold DA, Baek SH, Schork NJ, Rose DW, Larsen DD, Sachs BD, et al. RORa coordinates reciprocal signaling in cerebellar development through sonic hedgehog and calcium-dependent pathways. Neuron. 2003;40(6):1119–31. PubMed PMID: 14687547.

32. Cho YW, Hong CJ, Hou A, Gent PM, Zhang K, Won KJ, et al. Zfp423 binds autoregulatory sites in p19 cell culture model. PloS One. 2013;8(6):e66514. Epub 2013/06/14. doi: 10.1371/journal.pone.0066514. PubMed PMID: 23762491; PubMed Central PMCID: PMC3675209.

33. Jacobsen PF, Jenkyn DJ, Papadimitriou JM. Establishment of a human medulloblastoma cell line and its heterotransplantation into nude mice. J Neuropathol Exp Neurol. 1985;44(5):472–85. Epub 1985/09/01. PubMed PMID: 2993532.

34. Peyrl A, Krapfenbauer K, Slavc I, Yang JW, Strobel T, Lubec G. Protein profiles of medulloblastoma cell lines DAOY and D283: identification of tumor-related proteins and principles. Proteomics. 2003;3(9):1781–800. Epub 2003/09/16. doi: 10.1002/pmic.200300460. PubMed PMID: 12973738.

35. Wang Y, Zhou Z, Walsh CT, McMahon AP. Selective translocation of intracellular Smoothened to the primary cilium in response to Hedgehog pathway modulation. Proc Natl Acad Sci U S A. 2009;106(8):2623–8. Epub 2009/02/07. doi: 10.1073/pnas.0812110106. PubMed PMID: 19196978; PubMed Central PMCID: PMC2650314.

36. Kuzhandaivel A, Schultz SW, Alkhori L, Alenius M. Cilia-mediated hedgehog signaling in Drosophila. Cell Rep. 2014;7(3):672–80. Epub 2014/04/29. doi: 10.1016/j.celrep.2014.03.052. PubMed PMID: 24768000.

37. Chizhikov VV, Davenport J, Zhang Q, Shih EK, Cabello OA, Fuchs JL, et al. Cilia proteins control cerebellar morphogenesis by promoting expansion of the granule progenitor pool. J Neurosci. 2007;27(36):9780–9. Epub 2007/09/07. doi: 10.1523/JNEUROSCI.5586-06.2007. PubMed PMID: 17804638.

38. Heinz S, Benner C, Spann N, Bertolino E, Lin YC, Laslo P, et al. Simple combinations of lineage-determining transcription factors prime cis-regulatory elements required for macrophage and B cell identities. Mol Cell. 2010;38(4):576–89. Epub 2010/06/02. doi: 10.1016/j.molcel.2010.05.004. PubMed PMID: 20513432; PubMed Central PMCID: PMC2898526.

39. Levin JZ, Yassour M, Adiconis X, Nusbaum C, Thompson DA, Friedman N, et al. Comprehensive comparative analysis of strand-specific RNA sequencing methods. Nat Methods. 2010;7(9):709–15. Epub 2010/08/17. doi: 10.1038/nmeth.1491. PubMed PMID: 20711195; PubMed Central PMCID: PMC3005310.

40. Mootha VK, Lindgren CM, Eriksson KF, Subramanian A, Sihag S, Lehar J, et al. PGC-1alpha-responsive genes involved in oxidative phosphorylation are coordinately downregulated in human diabetes. Nat Genet. 2003;34(3):267–73. Epub 2003/06/17. doi: 10.1038/ng1180. PubMed PMID: 12808457.

41. Subramanian A, Tamayo P, Mootha VK, Mukherjee S, Ebert BL, Gillette MA, et al. Gene set enrichment analysis: a knowledge-based approach for interpreting genome-wide expression profiles. Proc Natl Acad Sci U S A. 2005;102(43):15545–50. Epub 2005/10/04. doi: 10.1073/pnas.0506580102. PubMed PMID: 16199517; PubMed Central PMCID: PMC1239896.

42. Wiederschain D, Chen L, Johnson B, Bettano K, Jackson D, Taraszka J, et al. Contribution of polycomb homologues Bmi-1 and Mel-18 to medulloblastoma pathogenesis. Mol Cell Biol. 2007;27(13):4968–79. Epub 2007/04/25. doi: 10.1128/MCB.02244-06. PubMed PMID: 17452456; PubMed Central PMCID: PMC1951487.

43. Concepcion D,Flores-Garcia L, Hamilton BA. Multipotent genetic suppression of retrotransposon-induced mutations by Nxf1 through fine-tuning of alternative splicing. PLoS Genet. 2009;5(5):e1000484. Epub 2009/05/14. doi: 10.1371/journal.pgen.1000484. PubMed PMID: 19436707; PubMed Central PMCID: PMC2674570.

44. Floyd JA, Gold DA, Concepcion D, Poon TH, Wang X, Keithley E, et al. A natural allele of Nxf1 suppresses retrovirus insertional mutations. Nat Genet. 2003;35(3):221–8. PubMed PMID: 14517553.

45. Badano JL, Mitsuma N, Beales PL, Katsanis N. The ciliopathies: an emerging class of human genetic disorders. Annu Rev Genomics Hum Genet. 2006;7:125–48. Epub 2006/05/26. doi: 10.1146/annurev.genom.7.080505.115610. PubMed PMID: 16722803.

46. Lee JE, Gleeson JG. Cilia in the nervous system: linking cilia function and neurodevelopmental disorders. Curr Opin Neurol. 2011;24(2):98–105. Epub 2011/03/10. doi: 10.1097/WCO.0b013e3283444d05. PubMed PMID: 21386674.

47. Louie CM, Gleeson JG. Genetic basis of Joubert syndrome and related disorders of cerebellar development. Hum Mol Genet. 2005;14 Spec No. 2:R235–42. Epub 2005/10/26. doi: 10.1093/hmg/ddi264. PubMed PMID: 16244321.

48. Wolf MT, Hildebrandt F. Nephronophthisis. Pediatr Nephrol. 2011;26(2):181–94. Epub 2010/07/24. doi: 10.1007/s00467-010-1585-z. PubMed PMID: 20652329.

49. Hu Q, Milenkovic L, Jin H, Scott MP, Nachury MV, Spiliotis ET, et al. A septin diffusion barrier at the base of the primary cilium maintains ciliary membrane protein distribution. Science. 2010;329(5990):436–9. Epub 2010/06/19. doi: 10.1126/science.1191054. PubMed PMID: 20558667; PubMed Central PMCID: PMC3092790.

50. Milenkovic L, Scott MP, Rohatgi R. Lateral transport of Smoothened from the plasma membrane to the membrane of the cilium. J Cell Biol. 2009;187(3):365–74. Epub 2009/12/02. doi: 10.1083/jcb.200907126. PubMed PMID: 19948480; PubMed Central PMCID: PMC2779247.

51. Piperno G, LeDizet M, Chang XJ. Microtubules containing acetylated alpha-tubulin in mammalian cells in culture. J Cell Biol. 1987;104(2):289–302. Epub 1987/02/01. PubMed PMID: 2879846; PubMed Central PMCID: PMC2114420.

52. Hubbert C, Guardiola A, Shao R, Kawaguchi Y, Ito A, Nixon A, et al. HDAC6 is a microtubule-associated deacetylase. Nature. 2002;417(6887):455–8. Epub 2002/05/25. doi: 10.1038/417455a. PubMed PMID: 12024216.

53. Liu A, Wang B, Niswander LA. Mouse intraflagellar transport proteins regulate both the activator and repressor functions of Gli transcription factors. Development. 2005;132(13):3103–11. Epub 2005/06/03. doi: 10.1242/dev.01894. PubMed PMID: 15930098.

54. May SR, Ashique AM, Karlen M, Wang B,Shen Y, Zarbalis K, et al. Loss of the retrograde motor for IFT disrupts localization of Smo to cilia and prevents the expression of both activator and repressor functions of Gli. Dev Biol. 2005;287(2):378–89. Epub 2005/10/19. doi: 10.1016/j.ydbio.2005.08.050. PubMed PMID: 16229832.

55. Caspary T, Larkins CE, Anderson KV. The graded response to Sonic Hedgehog depends on cilia architecture. Dev Cell. 2007;12(5):767–78. Epub 2007/05/10. doi: 10.1016/j.devcel.2007.03.004. PubMed PMID: 17488627.

56. Pazour GJ, Dickert BL, Vucica Y, Seeley ES, Rosenbaum JL, Witman GB, et al. Chlamydomonas IFT88 and its mouse homologue, polycystic kidney disease gene tg737, are required for assembly of cilia and flagella. J Cell Biol. 2000;151(3):709–18. Epub 2000/11/04. PubMed PMID: 11062270; PubMed Central PMCID: PMC2185580.

57. Taulman PD, Haycraft CJ, Balkovetz DF, Yoder BK. Polaris, a protein involved in left-right axis patterning, localizes to basal bodies and cilia. Mol Biol Cell. 2001;12(3):589–99. Epub 2001/03/17. PubMed PMID: 11251073; PubMed Central PMCID: PMC30966.

58. Cameron DA, Pennimpede T, Petkovich M. Tulp3 is a critical repressor of mouse hedgehog signaling. Dev Dyn. 2009;238(5):1140–9. Epub 2009/04/01. doi: 10.1002/dvdy.21926. PubMed PMID: 19334287.

59. Norman RX, Ko HW, Huang V, Eun CM, Abler LL, Zhang Z, et al. Tubby-like protein 3 (TULP3) regulates patterning in the mouse embryo through inhibition of Hedgehog signaling. Hum Mol Genet. 2009;18(10):1740–54. Epub 2009/03/17. doi: 10.1093/hmg/ddp113. PubMed PMID: 19286674; PubMed Central PMCID: PMC2671991.

60. Patterson VL, Damrau C, Paudyal A, Reeve B, Grimes DT, Stewart ME, et al. Mouse hitchhiker mutants have spina bifida, dorso-ventral patterning defects and polydactyly: identification of Tulp3 as a novel negative regulator of the Sonic hedgehog pathway. Hum Mol Genet. 2009;18(10):1719–39. Epub 2009/02/19. doi: 10.1093/hmg/ddp075. PubMed PMID: 19223390; PubMed Central PMCID: PMC2671985.

61. Mukhopadhyay S, Wen X, Chih B, Nelson CD, Lane WS, Scales SJ, et al. TULP3 bridges the IFT-A complex and membrane phosphoinositides to promote trafficking of G protein-coupled receptors into primary cilia. Genes Dev. 2010;24(19):2180–93. Epub 2010/10/05. doi: 10.1101/gad.1966210. PubMed PMID: 20889716; PubMed Central PMCID: PMC2947770.

62. Mukhopadhyay S, Wen X, Ratti N, Loktev A, Rangell L, Scales SJ, et al. The ciliary G-protein-coupled receptor Gpr161 negatively regulates the Sonic hedgehog pathway via cAMP signaling. Cell. 2013;152(1-2):210–23. Epub 2013/01/22. doi: 10.1016/j.cell.2012.12.026. PubMed PMID: 23332756.

63. Qin J, Lin Y, Norman RX, Ko HW, Eggenschwiler JT. Intraflagellar transport protein 122 antagonizes Sonic Hedgehog signaling and controls ciliary localization of pathway components. Proc Natl Acad Sci U S A. 2011;108(4):1456–61. Epub 2011/01/07. doi: 10.1073/pnas.1011410108. PubMed PMID: 21209331; PubMed Central PMCID: PMC3029728.

64. Jung B, Messias AC, Schorpp K, Geerlof A, Schneider G, Saur D, et al. Novel small molecules targeting ciliary transport of Smoothened and oncogenic Hedgehog pathway activation. Sci Rep. 2016;6:22540. Epub 2016/03/05. doi: 10.1038/srep22540. PubMed PMID: 26931153; PubMed Central PMCID: PMC4773810.

65. Jung B, Padula D, Burtscher I, Landerer C, Lutter D, Theis F, et al. Pitchfork and Gprasp2 Target Smoothened to the Primary Cilium for Hedgehog Pathway Activation. PLoS One. 2016;11(2):e0149477. Epub 2016/02/24. doi: 10.1371/journal.pone.0149477. PubMed PMID: 26901434; PubMed Central PMCID: PMC4763541.

66. Thomas J, Morle L, Soulavie F, Laurencon A, Sagnol S, Durand B. Transcriptional control of genes involved in ciliogenesis: a first step in making cilia. Biol Cell. 2010;102(9):499–513. Epub 2010/08/10. doi: 10.1042/BC20100035. PubMed PMID: 20690903.

67. Choksi SP, Lauter G, Swoboda P, Roy S. Switching on cilia: transcriptional networks regulating ciliogenesis. Development. 2014;141(7):1427–41. Epub 2014/03/20. doi: 10.1242/dev.074666. PubMed PMID: 24644260.

68. Swoboda P, Adler HT, Thomas JH. The RFX-type transcription factor DAF-19 regulates sensory neuron cilium formation in C. elegans. Mol Cell. 2000;5(3):411–21. Epub 2000/07/06. PubMed PMID: 10882127.

69. Dubruille R, Laurencon A, Vandaele C, Shishido E, Coulon-Bublex M, Swoboda P, et al. Drosophila regulatory factor X is necessary for ciliated sensory neuron differentiation. Development. 2002;129(23):5487–98. Epub 2002/10/31. PubMed PMID: 12403718.

70. Bonnafe E, Touka M, AitLounis A, Baas D, Barras E, Ucla C, et al. The transcription factor RFX3 directs nodal cilium development and left-right asymmetry specification. Mol Cell Biol. 2004;24(10):4417–27. Epub 2004/05/04. PubMed PMID: 15121860; PubMed Central PMCID: PMC400456.

71. Ashique AM, Choe Y, Karlen M, May SR, Phamluong K, Solloway MJ, et al. The Rfx4 transcription factor modulates Shh signaling by regional control of ciliogenesis. Sci Signal. 2009;2(95):ra70. Epub 2009/11/06. doi: 10.1126/scisignal.2000602. PubMed PMID: 19887680.

72. Piasecki BP, Burghoorn J, Swoboda P. Regulatory Factor X (RFX)-mediated transcriptional rewiring of ciliary genes in animals. Proc Natl Acad Sci U S A. 2010;107(29):12969–74. Epub 2010/07/10. doi: 10.1073/pnas.0914241107. PubMed PMID: 20615967; PubMed Central PMCID: PMC2919930.

73. Brody SL, Yan XH, Wuerffel MK, Song SK, Shapiro SD. Ciliogenesis and left-right axis defects in forkhead factor HFH-4-null mice. Am J Respir Cell Mol Biol. 2000;23(1):45–51. Epub 2000/06/29. doi: 10.1165/ajrcmb.23.1.4070. PubMed PMID: 10873152.

74. Stubbs JL, Oishi I, Izpisua Belmonte JC, Kintner C. The forkhead protein Foxj1 specifies node-like cilia in Xenopus and zebrafish embryos. Nat Genet. 2008;40(12):1454–60. Epub 2008/11/18. doi: 10.1038/ng.267. PubMed PMID: 19011629; PubMed Central PMCID: PMC4648715.

75. Yu X, Ng CP, Habacher H, Roy S. Foxj1 transcription factors are master regulators of the motile ciliogenic program. Nat Genet. 2008;40(12):1445–53. Epub 2008/11/18. doi: 10.1038/ng.263. PubMed PMID: 19011630.

76. Cruz C, Ribes V, Kutejova E, Cayuso J, Lawson V, Norris D, et al. Foxj1 regulates floor plate cilia architecture and modifies the response of cells to sonic hedgehog signalling. Development. 2010;137(24):4271–82. Epub 2010/11/26. doi: 10.1242/dev.051714. PubMed PMID: 21098568; PubMed Central PMCID: PMC2990214.

77. Lu H, Toh MT, Narasimhan V, Thamilselvam SK, Choksi SP, Roy S. A function for the Joubert syndrome protein Arl13b in ciliary membrane extension and ciliary length regulation. Dev Biol. 2015;397(2):225–36. Epub 2014/12/03. doi: 10.1016/j.ydbio.2014.11.009. PubMed PMID: 25448689.

78. Kile BT, Hentges KE, Clark AT, Nakamura H, Salinger AP, Liu B, et al. Functional genetic analysis of mouse chromosome 11. Nature. 2003;425(6953):81–6. PubMed PMID: 12955145.

79. Hatten ME. Neuronal regulation of astroglial morphology and proliferation in vitro. J Cell Biol. 1985;100(2):384–96. Epub 1985/02/01. PubMed PMID: 3881455; PubMed Central PMCID: PMC2113456.

80. Hatten ME, Gao W-Q, Morrison ME, Mason CA. The Cerebellum: Purification and Coculture of Identified Cell Populations. In: Banker G, Goslin K, editors. Culturing Nerve Cells. 2nd ed. ed. Cambridge, MA: MIT Press; 1998.

81. Chen JK, Taipale J, Cooper MK,Beachy PA. Inhibition of Hedgehog signaling by direct binding of cyclopamine to Smoothened. Genes Dev. 2002;16(21):2743–8. Epub 2002/11/05. doi: 10.1101/gad.1025302. PubMed PMID: 12414725; PubMed Central PMCID: PMC187469.

82. Stewart SA, Dykxhoorn DM, Palliser D, Mizuno H, Yu EY, An DS, et al. Lentivirus-delivered stable gene silencing by RNAi in primary cells. RNA. 2003;9(4):493–501. Epub 2003/03/22. PubMed PMID: 12649500; PubMed Central PMCID: PMC1370415.

83. DobinA, DavisCA, SchlesingerF, Drenkow J, Zaleski C, Jha S, et al. STAR: ultrafast universal RNA-seq aligner. Bioinformatics.2013;29(1):15–21.Epub 2012/10/30. doi: 10.1093/bioinformatics/bts635. PubMed PMID: 23104886; PubMed Central PMCID: PMC3530905.

84. Anders S, McCarthy DJ, Chen Y, Okoniewski M, Smyth GK, Huber W, et al. Count-based differential expression analysis of RNA sequencing data using R and Bioconductor. Nat Protoc. 2013;8(9):1765–86. Epub 2013/08/27. doi: 10.1038/nprot.2013.099. PubMed PMID: 23975260.

85. Robinson MD, McCarthy DJ, Smyth GK. edgeR: a Bioconductor package for differential expression analysis of digital gene expression data. Bioinformatics. 2010;26(1):139–40. Epub 2009/11/17. doi: 10.1093/bioinformatics/btp616. PubMed PMID: 19910308; PubMed Central PMCID: PMC2796818.

